# Multiomic analysis identifies suppressive myeloid cell populations in human TB granulomas

**DOI:** 10.1101/2025.03.10.642376

**Authors:** Neharika Jain, Emmanuel C. Ogbonna, Zoltan Maliga, Connor Jacobson, Liang Zhang, Angela Shih, Jacob Rosenberg, Haroon Kalam, Andréanne Gagné, Isaac H. Solomon, Sandro Santagata, Peter K. Sorger, Bree B. Aldridge, Amanda J. Martinot

## Abstract

Tuberculosis (TB) remains a major global health challenge, particularly in the context of multidrug-resistant (MDR) *Mycobacterium tuberculosis* (Mtb). Host-directed therapies (HDTs) have been proposed as adjunctive therapy to enhance immune control of infection. Recently, one such HDT, pharmacologic modulation of myeloid-derived suppressor cells (MDSCs), has been proposed to treat MDR-TB. While MDSCs have been well characterized in cancer, their role in TB pathogenesis remains unclear. To investigate whether MDSCs or other myeloid suppressor populations contribute to TB granuloma microenvironments (GME), we performed spatial transcriptional profiling and single-cell immunophenotyping on eighty-four granulomas in lung specimens from three individuals with active disease. Granulomas were histologically classified based on H&E staining, and transcriptional signatures were compared across regions of interest (ROIs) at different states of granuloma maturation. Our analysis revealed that immune suppression within granuloma was not primarily driven by classical MDSCs but rather by multiple myeloid cell subsets, including dendritic cells expressing indoleamine 2,3 dioxygenase-1 expressing (IDO1+ DCs). IDO1+ DCs were the most frequently observed suppressive myeloid cells, particularly in cellular regions, and their spatial proximity to activated T cells suggested localized immunosuppression. Importantly, granulomas at different stages contained distinct proportions of suppressor myeloid cells, with necrotic and cellular regions showing different myeloid phenotypes that may influence granuloma progression. Gene set enrichment analysis (GSEA) further indicated that elevated IDO1 expression was associated with a complex immune response that balanced suppressive signaling, immune activation, and cellular metabolism. These findings suggest that classical MDSCs, as defined in tumor microenvironments, likely play a minor role in TB, whereas IDO1+ DCs may be key regulators of immune suppression in granulomas influencing local *Mtb* control in infected lung. A deeper understanding of the role of IDO1+ suppressive myeloid cells in TB granulomas is essential to assessing their potential as therapeutic targets in TB treatment.

## Introduction

TB remains one of the major causes of death world-wide. HIV-infected individuals are particularly susceptible, with ¼ of HIV infected individuals dying from TB disease. Even those on systemic antiretroviral medications remain hypersusceptible to tuberculosis (TB) disease and have one of the highest rates of MDR TB world-wide. Other co-morbidities have been linked to TB susceptibility; these include diabetes and metabolic disease. Moreover, even non-immune compromised individuals are also at higher risk of encountering MDR TB in endemic areas. The only approved vaccine for TB, BCG, is ineffective in preventing pulmonary TB in adults. Hence, there is continued interest in identifying and prioritizing new treatment modalities for drug-resistant TB in both immune suppressed and non-suppressed populations. Pharmacologic modulation of MDSC has been extensively studied in oncology and has been proposed as a potential HDT for use in TB patients as an adjunctive treatment to antimicrobial therapy.^1^ In cancer, MDSCs have been associated with immune-suppression and dampening of the host’s immune response locally in tissue microenvironments, but the role of MDSCs in *Mycobacterium tuberculosis (Mtb)* disease progression is unclear.

MDSCs share many features in common with both neutrophils and macrophages. By surface marker expression or histopathology, they are very similar to other mature and immature myeloid populations. Thus, a multi-marker approach is necessary to distinguish MDSCs from other myeloid populations. Human MDSCs have been grouped as polymorphonuclear (PMN)-MDSCs and monocytic (M)-MDSCs. ^2^ In tumors, they comprise a distinct part of the tumor microenvironment and their presence in tumors is linked with decreased cytotoxic T cell function and tumor progression^3–5^. However, less is known about the role of MDSCs in infectious disease, such as TB or HIV, where both detrimental and beneficial effects have been reported.^6–8^In TB, the few reports investigating MDSCs in people have associated their presence with disease progression.^9^ It has been shown that MDSCs accumulate in the blood and pleural fluid during active TB disease, and that circulating numbers of MDSCs decrease after successful chemotherapy. However, it is unknown whether MDSCs traffic to granulomas and whether and to what extent they contribute to granuloma structure and function *in vivo* in TB patients. *In vitro* evidence suggests that MDSCs phagocytose but do not kill *Mtb* and may contribute to bacterial persistence both by harboring bacilli and interfering with the effectiveness of macrophage anti-microbial responses.^10^ Recent studies in non-human primates have suggested that MDSCs play important roles in the immunoregulation in and around TB granulomas. ^11^ IDO1 expression is high in human TB granulomas ^12^ and in the rhesus macaques, the administration of the IDO1 inhibitor D-1MT lead to significantly lower bacterial burden, increased T cell recruitment and better lung pathology with lower involvement in comparison to control infected (non-treated) animals.

A better understanding of MDSCs phenotypes in *Mtb* infected tissues during active TB disease is essential before exploring MDSCs modulation as a HDT for TB. We used two complimentary technologies – spatial transcriptional profiling using GeoMx (Bruker)^13^ and tissue-cyclic immunofluorescence (t-CyCIF)^14^ – to characterize myeloid cell populations and their relationship with TB granulomas across various stages of development and disease severity. Our findings reveal that traditionally defined MDSCs are present in TB-infected lungs and are associated with both active and quiescent granulomas, though they represent only a small fraction of the suppressive myeloid cells within these structures. The phenotype of suppressive myeloid cells varies depending on the stage of granuloma development. Notably, a highly suppressive granuloma microenvironment (GME) is more prevalent in cellular lesions lacking necrosis. We also observed that *ido1* expression by suppressive myeloid cells within granuloma is paradoxically associated with increased regulatory T cell activation and the absence of granuloma necrosis. Based on these findings, we propose that IDO expression by suppressive myeloid cells is critical in resolving tissue damage and control leading *Mtb* within granulomas. Our data suggest that a deeper understanding of how IDO1 regulates lung immunity during TB infection is necessary before pursuing pharmacologic modulation of suppressive myeloid cells as HDT for TB.

## Results

### Interindividual and intraindividual granuloma heterogeneity exists in human TB infected lung

Formalin-fixed paraffin-embedded (FFPE) tissues were selected from archived biopsies and autopsies processed between 1990 and 2018 at Massachusetts General Hospital and Brigham and Women’s Hospital in Boston, MA in which tissue-based examination of the lung or thoracic lymph node indicated granulomatous inflammation as the primary process, with mycobacterial infection in the differential diagnosis (88 cases). Two pulmonary pathologists evaluated the samples using clinical histopathology techniques, including H&E and Ziehl-Neelsen acid-fast staining. After reviewing case files, 15 cases were determined to have clinical and microbiologic confirmation/high suspicion of TB infection of which three cases, from one female and two males (referred to as case 2, 3, and 4 **(Suppl Table 1)**.

Specimens exhibited a range of multiple granuloma phenotypes: (i) ranging from necrotizing/active granulomas, marked by central caseous necrosis surrounded by immune cells, (ii), cellular (inactive/non-necrotizing) granulomas, lacking necrosis), and consisting mainly of epithelioid macrophages and lymphocytes, and (iii) cellular granulomas, characterized by dense aggregates of macrophages, T lymphocytes, and abundant collagen deposition indicating a resolved state. These histopathologic features varied across the selected cases, which reflected differences in their clinical presentations and medical histories (**Fig.1A-F**, **Suppl. Table 1**). Case 3 presented the most diverse morphology, featuring numerous necrotizing granulomas along with cellular and fibrotic granulomas, as well as prominent tertiary lymphoid structures (TLS) interspersed among the granulomas (**Fig.1A, D**). Case 2 exhibited predominantly, large caseating granulomas (**Fig.1B, E**), while Case 4 showed few necrotic granulomas but a predominance of cellular granulomas (**Fig. 1C, F**).

**Fig. 1.**
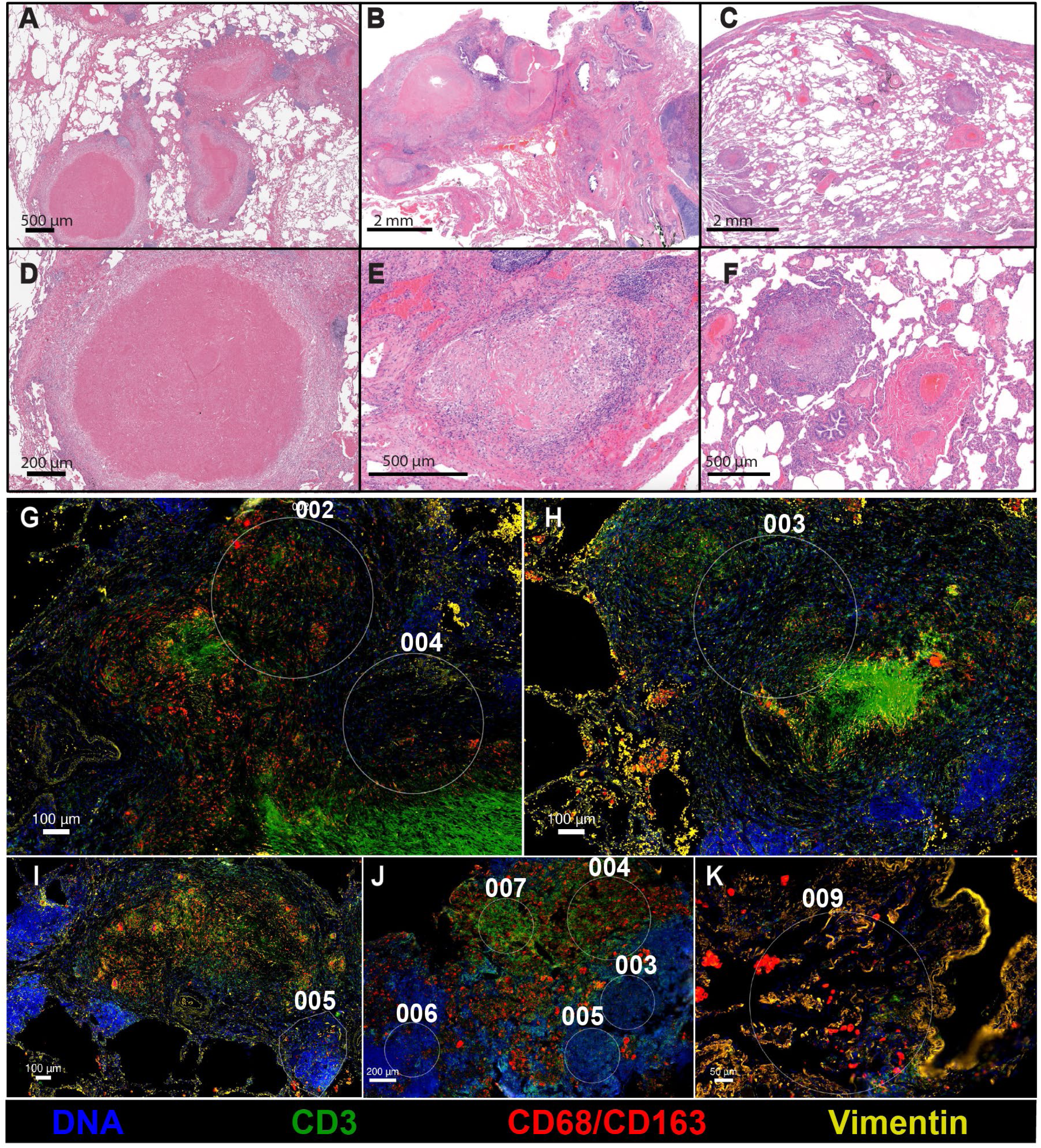
Heterogeneous human granuloma subtypes were selected and profiled using spatial transcriptomics. Representative H&E images of granulomas from **A)** Case 3, **B)** Case 2, and **C)** Case 4 showing a range of granuloma morphology with **D)** necrotizing granulomas, **E)** cellular non-necrotizing granulomas, and **F)** small, lymphocyte rich granulomas as assessed by a pulmonary pathologist using traditional morphological annotation. Regions of interest (ROIs) (**G-K**) were selected based on morphological and fluorescence staining for DNA (blue), CD3 (green), CD68/CD163 (red), and vimentin (yellow). ROIs bordering necrotic **(G)** granulomas and **(H)** solid or cellular granulomas were selected in addition to tertiary lymphoid structures **(I),** lymph node **(J),** and uninvolved lung **(K)** for spatial transcriptional profiling using the human whole transcriptome probe set using GeoMx (Bruker).

We performed microregional transcriptional profiling and multiplexed tissue immunofluorescence on serial sections to investigate whether MDSCs could be identified around active human TB granulomas. Our hypothesis was that heterogeneity in granuloma maturation states would correlate with variation in granuloma-associated microenvironments (GME), defined by regional immune cell infiltrates, including CD4^+^ and CD8^+^ T lymphocytes, B cells, and myeloid cells. We were particularly interested in determining whether the spatial localization of MDSCs varied across granuloma phenotypes. Given that MDSCs are known to suppress cytotoxic T lymphocyte function in tumors, we predicted that, if present in TB-infected lung, they would be associated with necrotic granulomas and linked to local T-cell dysfunction. Due to the limited data on the functional role and significance of MDSCs in TB infected lung tissues, we used an unbiased spatial whole transcriptome analysis (WTA) platform (GeoMx, Bruker, Inc) integrated with single-cell immunophenotyping via highly multiplexed tissue cyclic immunofluorescence (CyCIF) to initiate this study of human TB.

### Heterogenous myeloid cell composition in histologically similar granulomas

Lung sections from each of the three cases underwent immunofluorescence staining for macrophages (CD163 and CD68 in the same channel), T lymphocytes (CD3), connective tissue (vimentin) and nuclei/DNA (DAPI) to guide ROI selection (**Fig.1G-K)**, followed by spatial transcriptional profiling with human whole transcriptome atlas gene probe set (HsWTA, Bruker). Based on IF and H&E staining in serial sections with pathologist guidance, 35 ROIs were marked for processing and collection. All ROIs were pathologist-annotated as bordering either necrotic granulomas (NG, **Fig.1G**) or cellular granulomas (CG, **Fig.1H**), which may represent either resolving or reactivating lesions, in addition to lymphoid structures of the lung such as tertiary lymphoid structures (TLS, **Fig. 1I**), lymph node (LN, **Fig.1J**) and uninvolved lung (UL, **Fig.1K**). ROIs were selected to represent both T-cell rich and/or macrophage rich regions based on fluorescent protein expression of pan T cell marker (CD3) and combined macrophage markers (CD68/CD163)(**Suppl. Fig. 1**). Spatial transcriptomic data from each ROI underwent quality control check and quantile (Q3) normalization and gene counts for *cd3* and *cd68* confirmed correct selection of T-cell rich versus macrophage regions **(Suppl. Fig. 2)** followed by comparative analysis of annotations. Despite similar H&E features of granuloma types across cases, transcriptional data had prominent intra-patient clustering of transcripts by principal component analysis (PCA) irrespective of spatial localization and low variance was noted between defined annotations (CG, NG, LN, TLS and UL) (**Fig. 2A**). However, unsupervised clustering of ROIs grouped all three uninvolved lung (UL) ROIs (one from each case) as distinct from granulomas ROIs which served as the reference control for subsequent analysis. Variability in transcriptional data across the different ROI annotations were also apparent in the global heatmap of 10,187 gene transcripts profiled (**Fig. 2B**). While CG and NG granuloma annotations had similar global transcriptional profiles, lymphoid structures (LN and TLS) had distinct transcriptional profiles. Within individual annotations, remarkable variability in transcriptomes was also noted between ROIs of the same histological annotation from the same patient. Not all histological features were sampled from each patient specimen, for instance lymph node and TLS were only profiled from Case 2 and Case 3 respectively.

**Fig. 2.**
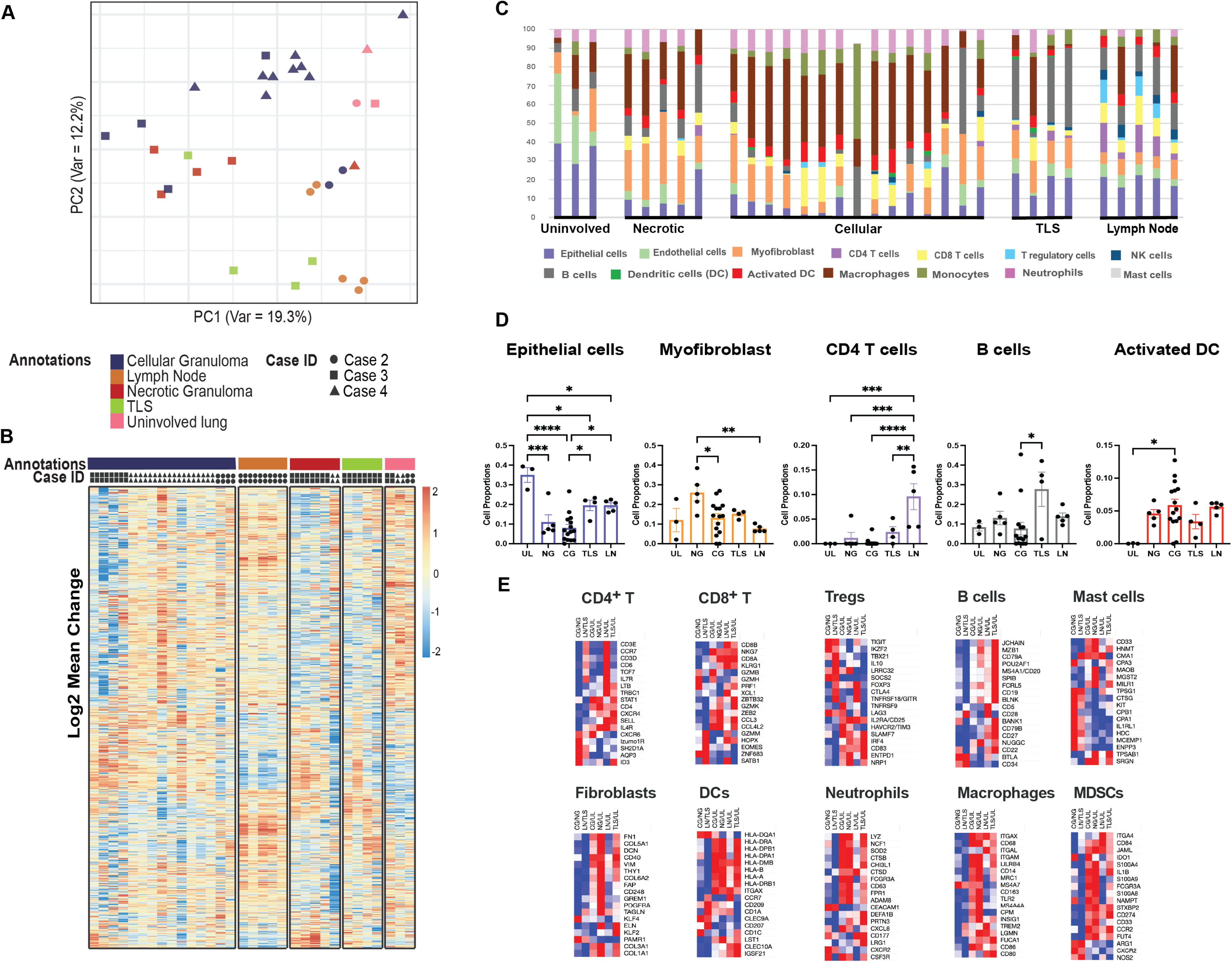
Spatial profiling of human TB granulomas reveals heterogenous composition of granuloma microenvironments across histologically similar granuloma subtypes. A) PCA plot showing variations among 5 different annotations assigned to all ROI across 3 cases. B) Supervised Clustering Heatmap depicting variations in assigned annotations from 3 different cases. C) Variability in Cellular deconvolution of individual ROI representing five different annotations. D) Bar graph representation of immune and non-immune cell types that show significant proportionality difference across annotations (Unpaired Mann-Whitney test; *, **, *** and **** represent p values ≤ 0.05, 0.01, 0.001 and 0.0001). E) Heatmap representing curated gene signature used in cell type calling for immune cell types.

Cellular granulomas (CG) ROIs had the best representation across all patients, with 15 ROIs in total from different cases. Necrotic granulomas were overrepresented in Case 3, limited in Case 4 and not sampled in Case 2, hence NG ROI sample size was small and biased towards Case 3.

Next, we used the transcriptional data to calculate the probabilistic cellular composition across individual ROIs using a cell deconvolution algorithm to approximate cell-type proportions based on single cell RNA seq (scRNAseq) reference data. (**Fig. 2C**). Currently, limited 10X scRNAseq data is available from human *Mtb* infected lung to generate cell deconvolution matrices for RNAseq analysis, hence, the healthy lung immune profile from Madissoon et. al. was adopted for reference in our study.^15,16^ Uninvolved lung ROIs, regions away from inflammation and immune infiltrates, were distinguished by alveolar epithelial cells (AT1), endothelial cells with matrix and stroma accounting for 60-80% of total cells. Cellular composition of granulomas ROIs were remarkably different from TLS and LN in percentage myeloid cells especially monocyte-macrophage populations and neutrophils. However, cellular granulomas were uniquely enriched in activated dendritic cells while necrotic granulomas had significantly higher proportion of myofibroblast cells.

Lymphocytic population comprising of CD4 T cells, T regulatory cells and NK cells had notable higher representation in lymph node ROIs while B cells were most prominent in TLS. We could not find any definitive pattern of CD8^+^ T cells as their distribution was quite variable across annotations (**Fig. 2D and Suppl Fig 3**). To support the cell type proportion model, we evaluated the relative expression of canonical genes associated with multiple cell types including B cells, CD4^+^ T lymphocytes, CD8^+^ T lymphocytes, T regulatory cells, neutrophils, macrophages, dendritic cells, mast cells and fibroblasts across each comparative annotation (**Fig. 2E**). Compared to uninvolved lung, TLS showed elevated expression of B cell genes, LN had higher expression of CD4^+^, CD8^+^ T lymphocytes transcripts, while cellular and necrotic granulomas displayed upregulated gene expression of neutrophils, macrophages, and dendritic cells validating ROI annotations. In addition, cellular granulomas had higher expression of mast cell genes compared to necrotic granulomas.

Because scRNAseq data on MDSCs were not available for human lung, we were unable to estimate the proportion of MDSCs across ROIs. However, we evaluated relative expression of a curated list of MDSC-associated genes compared to uninvolved lung, many of which were elevated in granulomas (**Fig. 2E**).

### Localized immune suppression is prominently associated with cellular granulomas

To understand the local cellular signaling pathways along with associated genes involved in regulating the unique granuloma niche, we evaluated differentially expressed genes (DEG) across annotations. Because of multiple comparison groups, we had a large representation of DEG i.e. 933 transcripts across annotations comparisons whose distribution pattern is shown as Venn diagram **(Fig. 3A**) and represented as a global heatmap (**Suppl. Fig. 4)**. Out of 933 genes, 130 genes were uniquely differentially expressed in necrotic granulomas and a relatively small number of 26 genes were uniquely differentially expressed in cellular granulomas compared to uninvolved lungs (**Suppl. Table 2).** As we were interested in identifying potential hotspots of MDSCs, we complied genes of myeloid lineage markers and suppressive genes that have been shown to be potential MDSCs markers and used the same cell type associated canonical gene lists represented in Fig 2E to evaluate the magnitude of their significant differential gene expression (greater than Log2 fold and adj p-value >0.05) across our comparisons in addition to B cells, T lymphocytes, myeloid cells, and mast cells (**Fig. 3B**). Similar to the trends observed for macrophages and neutrophils, MDSC associated genes were also upregulated in granulomatous regions (CG and NG). Strikingly, some of the key suppressive genes ascribed to MDSC suppressive function including *ido1, nampt,*^17,18^ *arg1, and nos2,* were highly enriched in cellular granulomas as compared to necrotic granulomas (**Fig. 3B**). Abundant B-cell signature genes were upregulated in TLS as compared to uninvolved lung (TLS/UL). Lymph nodes compared to uninvolved lung (LN/UL) had significant expression of genes associated with CD4^+^ and CD8^+^ T lymphocytes including TCF7, CD79A, SELL, IL7R, TRBC1. A significant proportion of myeloid (*cd14*, *lilrb4*, *cd68, itgam*/*cd11b*, *sod2*) and neutrophil gene transcripts (*ncf1, lyz, ctsb*) were differentially expressed in both cellular and necrotic granulomas. However, necrotic granulomas overall had more diverse expression of key B cell transcripts *mzb1*, *cd79a* and T cell transcripts *cd4, sell, stat1* signifying extensive immune infiltration in that region. While we initially performed comparative analysis based on uninvolved lungs as the reference control, we were curious to see whether significant differences in expression pattern between cellular granuloma and necrotic granuloma might emerge to explain the local immune environment as restrictive or permissive respectively (CG/NG). We observed over-expression of *ido1*, a highly immunosuppressive gene in all the TB granulomas which is consistent with recent studies in TB^12,19–21^, but it was significantly differentially upregulated in cellular granulomas in comparison to necrotic granuloma marking it as an important distinguishing feature of non-necrotic granulomas (**Fig. 3B**). In addition, there was elevated *hla-dqa1* in cellular compared to necrotic granulomas, suggesting induction of MHC Class II antigen presentation in cellular, non-necrotic granulomas.

**Figure 3.**
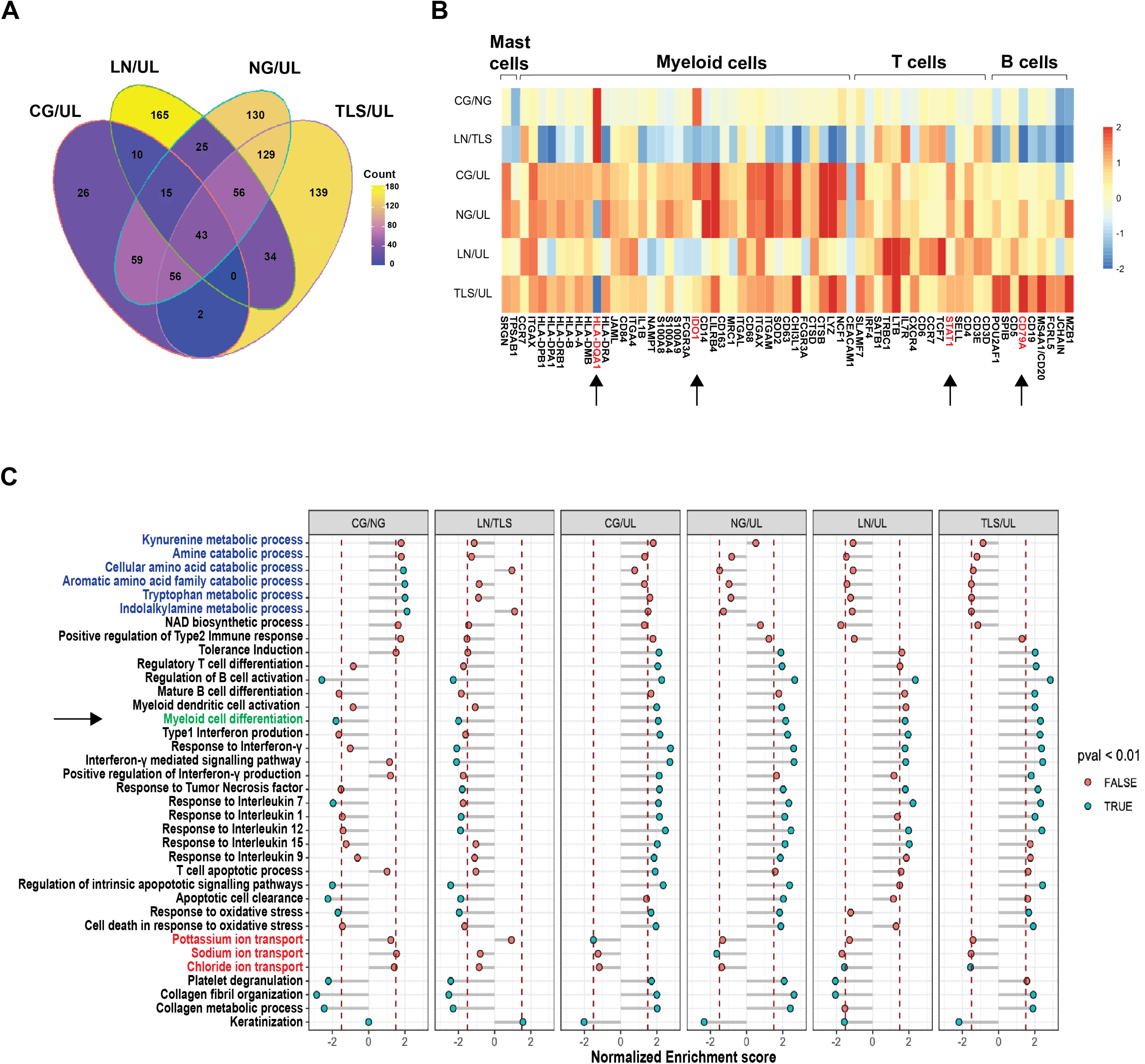
Cellular granulomas are defined by upregulation of suppression-related genes. **A)** Venn diagram highlighting the unique number of differentially expressed genes in defined annotations and varying degree of overlapping DEGs among the annotations **B)** Subset of DEG of important immune cells signature genes. DEGs (Differentially expressed genes) obtained from DESeq2 R-based pipeline in annotations (CG, NG, LN, TLS) compared to UL combined in unsupervised clustering manner to represent a heatmap. **C)** List of differential pathways significantly regulated in spatially defined annotations compared to UL or CG vs NG and LN vs TLS using Gene set enrichment analysis (GSEA). CG=cellular granuloma; NG=necrotic granuloma; LN=lymph node; TLS=tertiary lymphoid structure; UL=uninvolved lung.

Since suppressor cells are known to correlate with differential T cell recruitment and function, we subsequently performed gene set enrichment analysis (GSEA) to understand unique transcriptional programs that are active locally around granulomas as compared to other regions of the TB infected lung. Both cellular and necrotic granulomas upregulate transcriptional programs of major cytokine pathways known to influence *Mtb* pathophysiology including interferon gamma production and signaling, TNF signaling and signaling response to IL-1, IL-7, IL-9, IL-12, and IL-15, consistent with an antimicrobial cellular response **(Fig. 3C**).

Among the downregulated responses, we observed significant downregulation of K^+^ ion and Cl^−^ ion transport in the CG and NG ROIs respectively, supporting potential *Mtb* growth inhibition and clearance in CG ROIs. As both inorganic ions (K+ and Cl-) are necessary for *Mtb* survival, K^+^ ion-channel blockers are proposed as potential anti-TB agent as they disrupt the oxidative and pH balance in the cells restricting *Mtb* growth.^22–24^ A striking difference between cellular granulomas and necrotic granulomas was the upregulation of pathways associated with IDO metabolism including tryptophan and kynurenine metabolic processes consistent with the elevated IDO transcript level noted in cellular granulomas. High co-expression of IDO associated metabolic pathways within cellular granulomas supported our hypothesis that suppressor cells may be differentially associated with TB granulomas in different maturation states. Notably, only trends in elevated immune tolerance and Type II immune responses were noted in cellular granulomas versus necrotic granulomas. Apoptotic signaling pathways, cell clearance, and oxidative stress signaling pathways were differentially diminished in cellular granulomas (**Fig. 3C**).

### TB granulomas are composed of phenotypically diverse suppressive myeloid cell populations

Next, we aimed to phenotype single cells in both necrotic and cellular granuloma lesions to better assign suppressive genes to specific cellular infiltrates. We performed highly multiplexed tissue cyclic immunofluorescence (t-CyCIF) in serial sections from specimens profiled for spatial transcriptomics using a 50+ antibody panel optimized for human samples (**Suppl. Table 3**). ^14,25,26^ In contrast to GeoMx which provided bulk-RNA analysis within individual ROIs, CyCIF enabled comprehensive assessment of single-cell phenotypes across the entire tissue specimen resulting in deep analysis and immunophenotype of every cell of the infected TB lung specimen. Again, with pathologist guidance, granulomas in each of the three patient specimens were annotated as either necrotic or cellular. In contrast to GeoMx, in which we were limited by number and size of ROI that could be profiled, using CyCIF, we were able to profile all visible granulomas across the entire patient sample to further enrich the dataset. Granulomas that were transitional, defined as containing small areas of acellular material surrounded by fibrosis, were combined with solid, cellular granulomas in the analysis. A total of 34 necrotic and 50 cellular granulomas were annotated across the three lung specimens **(Figs. 1, 4A)**. A combination of surface, intracellular and nuclear markers were used and individually thresholded using an open-source visual gating tool (https://github.com/labsyspharm/minerva_analysis). Cell type calling was defined based on gating strategies adapted from flow cytometry and microscopic panels used for identifying multiple immune and non-immune cell types including MDSC (**Suppl. Fig. 5**). We validated cell-type calling by analyzing specific marker expression in different cell types within the three tissues (**Fig. 4B**).

**Figure 4:**
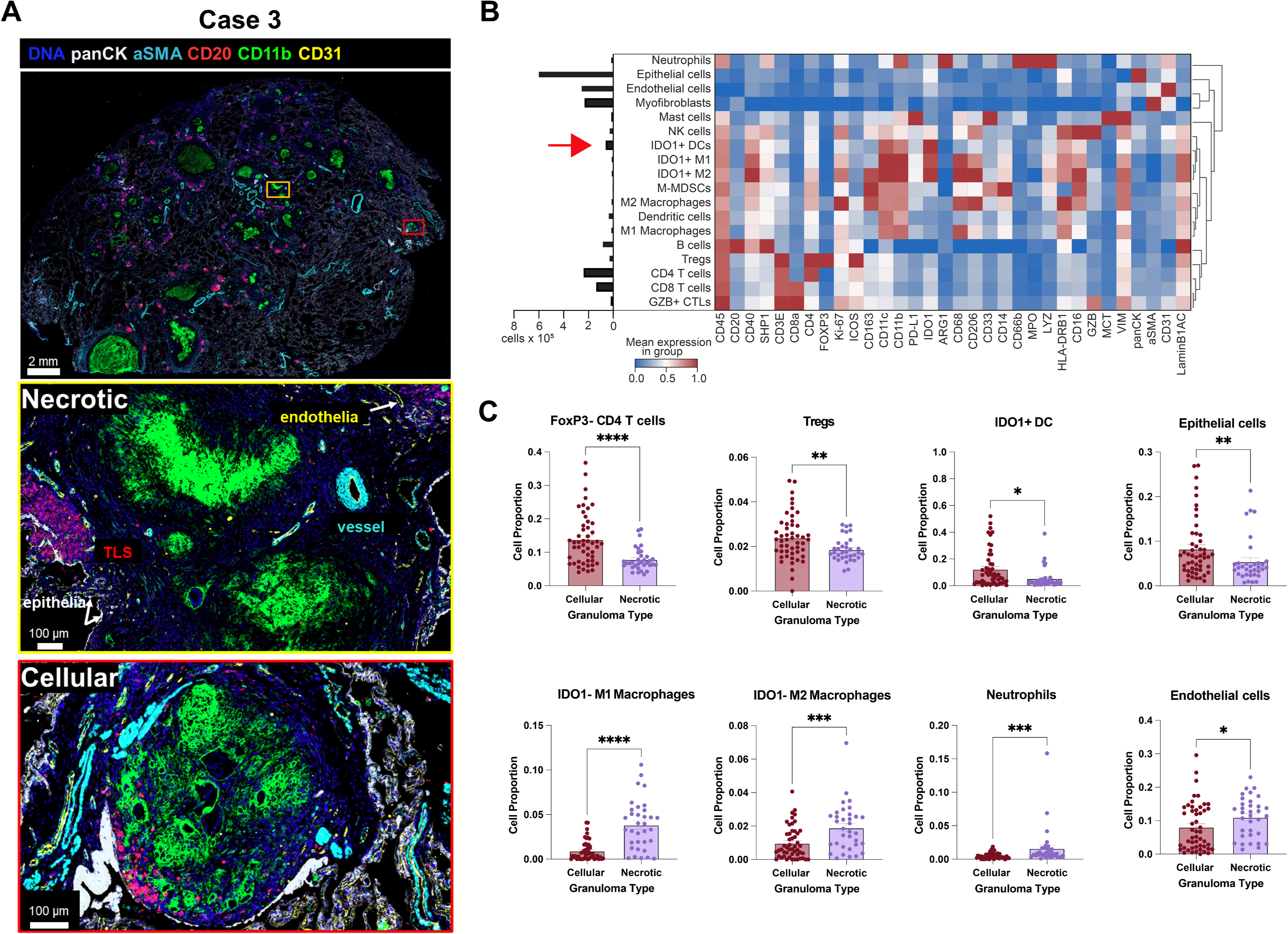
C**y**clic **immunofluorescence (CyCIF) imaging of human TB-infected lung samples reveals distinct suppressive cell populations. A)** TB-infected samples were imaged with CyCIF. Representative whole –slide image show tissue architectural markers – panCK (epithelia), aSMA (myofibroblasts), CD31 (endothelia), CD11b+ myeloid cells and CD20+ B-cells). Inset images highlight fields of view with variable granuloma types. **B)** Heatmap showing mean expression levels of protein markers in cell types. Dendrogram shows hierarchical/agglomerative clustering of cell phenotypes (Euclidean distance, complete linkage). Phenotyping was performed on cells as described in **Supplemental Fig. 4**. **C)** Comparative phenotypic analysis of necrotic and non-necrotic granulomas. Proportions of cell phenotypes are averaged across the granulomas of the same type. Error bars show standard error of mean. Unpaired Mann-Whitney test was used to assess significant differences in averaged proportions between granuloma types. p values > 0.05 (ns) not significant; *, **, *** and **** represent P values ≤ 0.05, 0.01, 0.001 and 0.0001, respectively. Plots of other phenotypes are shown in **Supplemental Fig. 5**.

Broadly, non-immune antibody markers include pan-cytokeratin (panCK), alpha smooth muscle actin (aSMA), and CD31 to stain the lung epithelia, airway smooth muscle, myofibroblasts, and endothelial cells, respectively. Similar to what we observed with GeoMx cell deconvolution, non-immune cells made up to 46% of the tissue cells with epithelial cells accounting for 25%, endothelial cells were 11% whereas myofibroblasts were 10% of all single cells. Immune cell types, distinguished by CD45 positivity were classified based on staining with one or more lineage-specific and relevant functional markers (**Suppl. Fig. 5**). CD3E, CD4, CD8a, FOXP3, and Granzyme B (GZB) were used in different combinations to identify T cell subsets including CD4^+^ helper T cells, regulatory T cells (Tregs), CD8^+^ cytotoxic T cells, and GZB^+^ cytotoxic T lymphocytes (GZB^+^ CTLs). Expression of other markers in T cell subsets was important in delineating their functions and capturing the spectrum of T-cells in the lung tissue. For example, there was a relatively higher expression of ICOS (a marker of activation) in FOXP3^+^ Tregs; and high Ki-67 expression in GZB^+^ cytotoxic T cells compared to other T-cells, suggesting increased proliferation. Common myeloid cell populations, an important branch of innate immune response, were defined as M1 macrophages, M2 macrophages, dendritic cells, and neutrophils. In addition to these, B cells, NK cells, Mast cells and MDSCs were also included in our gating strategy. Classical MDSCs were classified as CD11b^+^CD33^+^ HLA-DR^−^ cells based on marker expression data in cancer immunology.^2^ They were further subclassified as M-MDSCs which were phenotyped as CD11b^+^CD33^+^HLA-DR^−^CD14^+^ cells and PMN-MDSCs as CD11b^+^CD33^+^HLA-DR^−^CD66b^+^ (**Suppl. Fig. 5**). Furthermore, a set of functional markers, IDO1, ARG1, PD-L1, Ki-67, and ICOS, were included with the surface markers for functional characterization of different phenotyped cells. As IDO1 is a major immunosuppressive marker and highly expressed gene around granulomas (**Fig. 3B**), we sub-phenotyped multiple IDO1+ myeloid cells as IDO+ MI macrophages, IDO+ M2 macrophages and IDO+ DCs. The resulting phenotypes IDO1+ M1(1.1%), IDO1+ M2 (1.1%), and IDO1+ DCs (2.5%) represented the majority of suppressive myeloid cells among which IDO1+ DCs were the most abundantly detected suppressive myeloid cell population (**Fig. 4B, arrow**). Other immune cells evaluated in TB infected lung included other innate immune cells, including mast cells (0.7%), neutrophils (1.3%), and natural killer (NK) cells (1.8%). Mast cells, identified by tryptase (MCT) expression depicted high CD33 and PD-L1 expression (76% and 100% positivity, respectively). Mast cells are known to be abundant in human lung and mucosa, contributing to fighting intracellular mycobacteria as a phagocyte or by releasing cytokines and chemokines.^27^ Neutrophils also express functional proteins including arginase (ARG1), myeloperoxidase (MPO), and lysozyme (LYZ), suggesting their roles in regulating inflammatory response and antimicrobial activity.^28–30^ NK cells (GZB^+^CD16^+^CD3E-) expressed Ki-67 and HLA-DRB1, highlighting their role in antigen presentation in addition to their killing capacity.^31^ Importantly, we identified only 0.2% of cells as M-MDSCs in the tissues and were unable to identify sufficient numbers of PMN-MDSCs (less than 40 cells were classified as PMN-like MDSCs), therefore PMN-MDSCs were excluded from down-stream analysis.

To assess the relationship between immune composition and granuloma histopathology, we evaluated the relative proportion of above-mentioned cell types within defined boundaries around granuloma subtypes – cellular granulomas (CG) and necrotic granulomas (NG). CD4^+^ T cells (FoxP3^−^CD4^+^), T regulatory cells (Tregs; FoxP3^+^CD4^+^), IDO1^+^ DCs, and epithelial cells were significantly enriched in cellular granulomas (Mann-Whitney, P < 0.05) while IDO1^−^ M1, IDO1^−^ M2, neutrophils and endothelial cells were enriched in necrotic granulomas (Mann-Whitney, P < 0.05) (**Fig. 4C**). The increased presence of IDO1^+^ DCs in cellular granulomas corresponded to observations from our GeoMx data. Classically defined M-MDSCs were equally present in both cellular and necrotic granulomas (**Suppl. Fig. 6**). Therefore, we concluded that transcriptional and protein signals associated with immunosuppression including IDO, are likely heavily derived from suppressive dendritic cells in cellular granulomas rather than MDSCs. Furthermore, suppressive myeloid cells were enriched in cellular granulomas rather than regions of necrosis as previously hypothesized.

### Spatial recurrent cellular neighborhoods in TB-infected lung are defining features of granuloma types

The presence of suppressor cells has been associated with decreased effector T cell function in the tumor microenvironment.^32^ Therefore, we wanted to better understand the granuloma microenvironment, cell-cell interactions, cell associations, and T cell composition of regions containing high numbers of suppressive myeloid cells. To interrogate spatial neighborhoods of immune and non-immune cells associated with the granulomas, we performed latent Dirichlet allocation algorithm (LDA) on the assigned cell phenotypes. With LDA, we identified seven granuloma microenvironments (GME1-GME7) each defined by the highest proportion of cell types among varying compositions of different immune and non-immune cells (**Fig. 5A**). GME1 contained the largest number of cells of all the microenvironments (∼35% of all cells in the dataset) and was largely composed of epithelial cells (**Fig. 5A, B**). We observed the presence of NK cells (3.6%), which are a major contributor to the innate immune response to TB infection.^33^ The second microenvironment, GME2, was composed of mostly CD3E^+^ T cells, which visually mapped to the lymphocytic cuff of granulomas (**Suppl. Fig. 7**). Classically defined M-MDSCs were most frequently identified within GME2 (**Fig. 5A**). These data align with previously published work identifying suppressive myeloid cells in the lymphocytic cuff of both human and non-human primate granulomas.^12,20^ However, in some granulomas, GME2-assigned cells were also associated with granulomas but not spatially restricted to the granuloma rim. GME3 corresponded mainly to the lung endothelium. Interestingly, over 50% of all assigned neutrophils were located in GME3, which also had the highest mean expression of CD66b, MPO, and ARG1. Alpha smooth muscle actin (aSMA)-positive myofibroblasts predominated GME4, which mostly physically mapped to blood vessels but also fibrosis around some granulomas (**Fig.5C**). GME5 appears as lymphoid aggregates rich in B cells, T cells and some IDO1^−^ dendritic cells, both granuloma-associated and randomly within the lung parenchyma as TLS. Importantly, two myeloid-rich neighborhoods, GME6 and GME7, were identified mostly within and around granulomas. Closer inspection of the marker expression pattern in these two neighborhoods revealed them to be consisting of functionally different myeloid populations. GME6 contained a high proportion of IDO1^+^ myeloid cells (dendritic cells and macrophages) and had the highest proportion of PD-L1^+^ cells and was therefore classified as an immunosuppressive GME. GME7 was also significantly enriched for macrophage and dendritic cell populations which lacked similar elevated IDO1 expression (**Fig. 5C**). In GME6, IDO1^+^ DCs are the most prevalent cell type (28%), followed by IDO1^+^ M1 (19%) and IDO1^+^ M2 macrophages (15%). Comparatively, IDO1^−^ DCs make up only 2.8% of ME6. These suggest that immunosuppressive myeloid populations are spatially localized in TB-infected lungs and play distinct roles in TB pathobiology. Notably, classically defined M-MDSCs were equally represented in both GME6 and GME7 and represented only a small proportion of immunosuppressive myeloid cells (0.13% and 0.17%, respectively) compared to IDO^+^ and IDO^−^ dendritic cells and macrophages.

**Figure 5:**
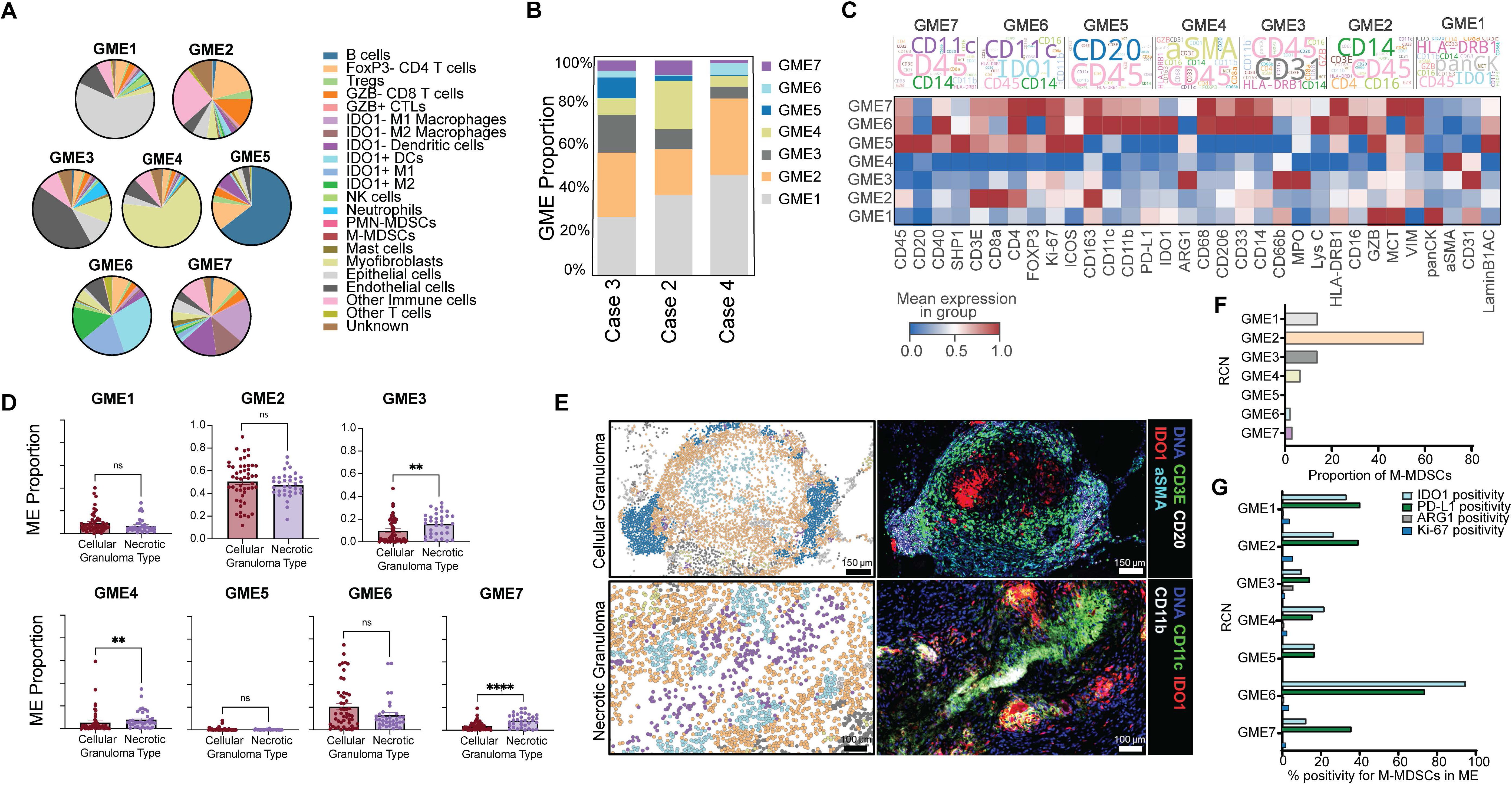
I**m**munoregulatory **spatial recurrent cellular neighborhoods are observed in TB-infected lung samples. A)** Spatial recurrent cellular neighborhoods (RCNs) or microenvironments (MEs) were computed using a k-nearest neighbor method based on the phenotype of each cell’s nearest neighbors (k=10). Pie charts show phenotypic composition of the 7 MEs. Based on the major cell types in each GME, a concise biological definition is thus: GME1, Epithelium; GME2, T-cell rich region; GME3, Endothelium; GME4, Fibroblast-rich; GME5, B-cell rich TLS; GME6, IDO1-high myeloid GME; GME7, IDO1-low myeloid GME. **B)** Composition of the GMEs in the different samples. **C)** Word cloud and heatmap showing protein marker expression levels in the GMEs. IDO1 (and PD-L1) expression distinguished the immunosuppressive GME6 from the other myeloid environment, GME7. In word cloud, only markers used for phenotyping are shown, with font size proportional to the intensity levels. **D)** Comparative analysis of necrotic and cellular (non-necrotic) granulomas showed significant differences in GME3, GME4 and GME7. Higher levels of GME6 in non-necrotic granulomas, although not significant (p value = 0.1829, two-tailed Mann-Whitney test; p value = 0.062, two-tailed t test). p values > 0.05 (ns). **E)** Representative CyCIF regions of interests (ROIs) highlighting GMEs in and around TB granulomas. For each FOV, in the first panel, the cells are colored based on the GMEs they belong to. Selected markers are shown depicting expression patterns in the GMEs. The top granuloma (cellular) had relatively high levels of GME6 corresponding to IDO1-expressing myeloid cells. In the bottom granuloma (necrotic), there appears to be a central GME7 region with IDO1-CD11c+ and CD11b+ cells with IDO1+ GME6 clusters nearby. The color scheme of GME corresponds to **Fig. 5B** legend. **F)** Proportion of M-MDSCs population in the various recurrent cellular networks (RCN) or granuloma microenvironments (GMEs). Monocytic-(M-) MDSCs localize in the T-lymphocyte-rich microenvironment GME2. **G)** Positivity of effector and proliferation protein markers, IDO1, PD-L1, ARG1, and Ki-67. GME2 has the highest population of M-MDSCs. However, the proportion of IDO1+ and PD-L1+ M-MDSCs to the MDSCs population in each GME was highest in GME6. The M-MDSCs in GME6 are almost all immunosuppressive.

To assess the relationship between granuloma histopathology and immune microenvironments, we compared the GME composition of necrotic granulomas versus cellular granulomas. GME3, GME4, and GME7 were overrepresented in necrotic granulomas (**Fig. 5D, E**). Higher numbers of neutrophils (as associated with GME3) were more prevalent in necrotic granulomas (**Fig. 4C**). Necrotic granuloma cores are typically rich in both viable and degenerative neutrophils.^34^ GME2 was not significantly different across granuloma subtypes.

GME6 was slightly enriched in cellular granulomas, however not significantly. The significant enrichment of GME7 in necrotic granulomas suggests that immunosuppression is not a feature of necrotic granulomas as compared to cellular granulomas, contrary to our original hypothesis (**Figs. 4C, 5D**).

Visual inspection of GME when spatially overlayed with CyCIF images suggested varying spatial arrangement of GME6 and GME7 in distinct granuloma subtypes and with other GMEs (**Fig. 5E**). For example, in the cellular granuloma depicted in **Fig. 5E** (**top**), GME6 (immunosuppressive with high IDO^+^ myeloid populations) was physically separated from GME7. In the necrotic granuloma shown, clusters of GME6 and GME7 were admixed within the granuloma (**Fig. 5E, bottom**). These data show that granulomas that differ by histologic features of necrosis have different granuloma microenvironments characterized by the presence or absence of unique suppressive myeloid cell subsets. In our data, both GME6 and GME7 represent microenvironments in close spatial proximity with the T-cell rich GME2 (**Fig. 5E, Suppl. Figs. 7B-C**). GME6 has the highest proportion of IDO^+^ DCs while M-MDSCs were overrepresented in GME2 associated with the lymphocytic cuff (**Fig. 5A, E, F**). Smaller numbers of MDSCs were identified in GME6 and GME7. We evaluated the proportion of suppressive “effector” molecules by CyCIF on these cells as a function of which microenvironment they were associated with to understand if their phenotype varied spatially. Relative to M-MDSCs population per GME, the positivity of key immunosuppressive markers IDO1 and PD-L1 was highest in ME6 – 95% and 74%, respectively (**Fig. 5G)**. These values were much lower in GME2 (27% IDO1^+^, 39% PD-L1^+^) and ME7 (12% IDO1^+^, 36% PD-L1^+^). Hence, while making up only a small proportion of the population of myeloid suppressor cells in granulomas, the M-MDSCs in ME6 are mostly immunosuppressive as defined by IDO1 positivity.

### IDO1^+^ DCs are enriched in cellular granulomas and have significant interactions with both classic and regulatory T lymphocytes

IDO1^+^ DCs are known to interact and recruit Tregs in cancer and have been associated with decreased cytotoxic T cell function in solid tumors. ^35,36^ We identified that IDO^+^ DCs are overrepresented in the immune suppressive granuloma microenvironment (GME6), whereas traditionally defined M-MDSCs were confined to GME2 representing the lymphocytic cuff of the majority of granulomas. Therefore, we wanted to understand how IDO1 positive and negative DCs and macrophages were interacting and affecting T lymphocytes in TB granulomas compared to classically defined M-MDSCs. To better characterize the spatial proximity of GME across different granuloma subsets we evaluated the spatial interactions of all immune cells with each other across TB infected lungs by a *k-*nearest-neighbor analysis (**Fig. 6**, **Suppl. Fig. 8**). IDO1+ DCs had significant interactions with T regulatory cells, CD4+ T cells, GZB+ CTLs, CD8 T cells, IDO1+ M1 and M2 macrophages across the lung samples (**Suppl. Fig. 8**). Given that the proportion of IDO1+ DCs differed across granuloma sub-types, we asked whether the spatial proximity of IDO1+ DCs differed across the different microenvironments. We scored the number of specific cell-to-cell interactions across all GME’s using spatial proximity volume (**Fig. 6A**). Spatial proximity volume is a spatial proximity metric defined as a ratio of the total number of cell interactions to the total number of cells in the dataset. IDO1^+^ DCs had the highest proximity volume to CD4^+^ and CD8^+^ T lymphocytes, regulatory T lymphocytes, and GZB^+^ CTLs in GME6 followed by GME2 (**Fig. 6A**). The spatial proximity of IDO1^+^ DCs in GME7 was much lower than that of GME6.

**Figure 6:**
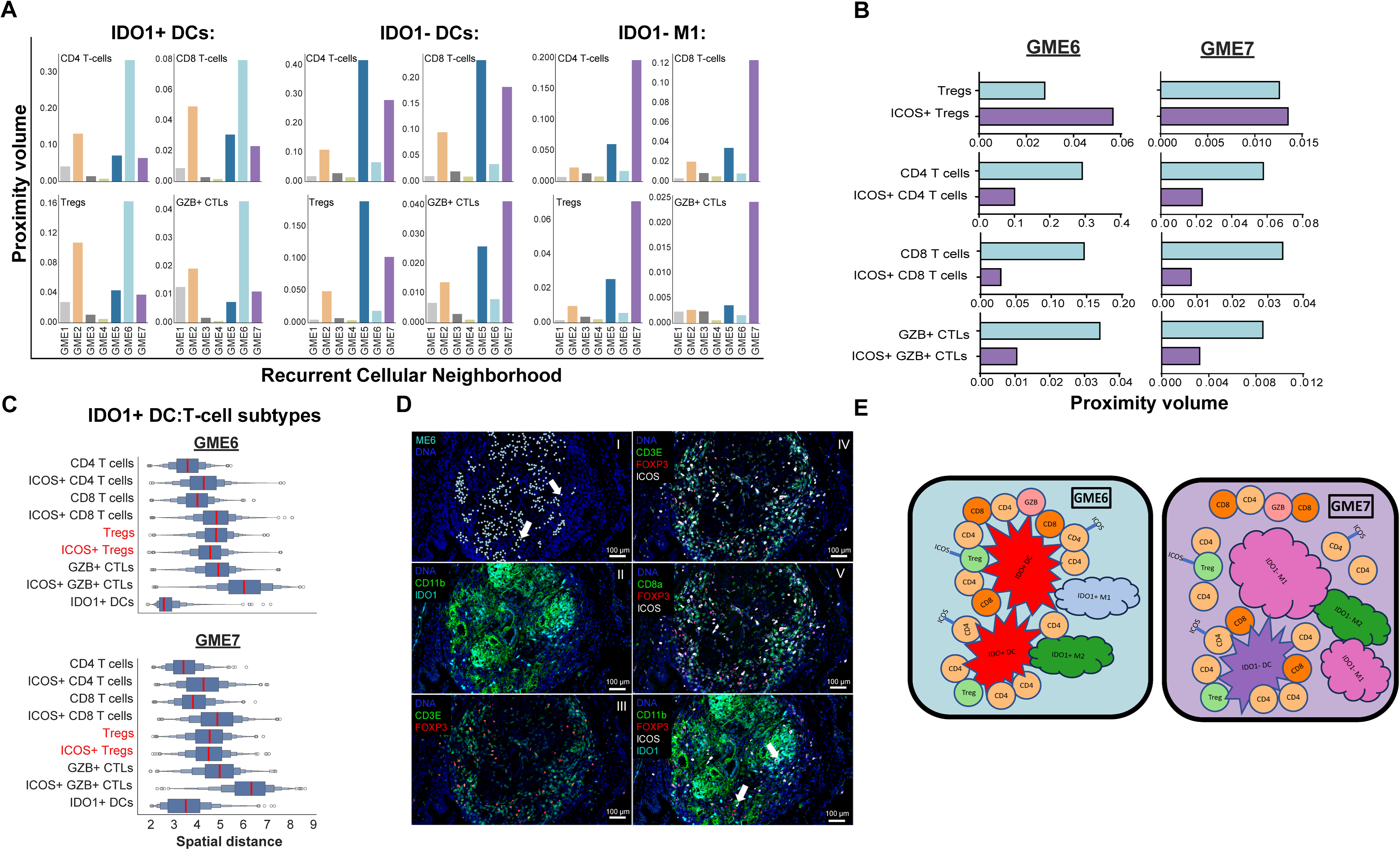
I**D**O1**+ dendritic cells drive cell-cell interactions within the immunosuppressive granuloma microenvironment. A)** Spatial proximity volume analysis scored the number of IDO1+ DCs, IDO1-DCs and IDO1-M1 macrophage interactions with T-cell subsets across the different GMEs. IDO1+ DCs drive the immunosuppression in GME6 which had the highest proximity volume between IDO1+ DCs and T cells. IDO1-DCs interacts with T-cells differentially: interacting with CD4 T-cells and Tregs highest in GME5 which corresponds to TLSs. IDO1-M1 macrophages make up the highest proportion of cells in GME7. These cells have the highest interactions with T-cells in GME7. **B)** Proximity volume scores between IDO1+ DCs and ICOS+ and ICOS-T cell subtypes in ME6 and ME7. In GME6, for most T cell subtypes except Tregs, ICOS+ (or activated) T cells had a lower spatial volume than ICOS-cells. This suggests that activated Tregs have more interactions to immunosuppressive IDO1+ DCs, while interactions between IDO1+ DCs and activated non-suppressive T cells are much less likely than their ICOS-counterparts. The specific interaction with activated Tregs was specific to GME6, not GME7. **C)** Spatial distance between T cells and IDO1+ DCs. In GME6, activated (ICOS+) Tregs are relatively closer to IDO1+ DCs and macrophages than ICOS-Tregs. This is unlike other T cell subtypes with ICOS+ subsets farther away. **D**) Representative images highlight the immunosuppressive GME6. I: Cells belonging to GME6 are colored cyan and overlaid onto DNA staining (DNA: blue). II: Significant expression of IDO1 by CD11b+ myeloid cells (DNA: blue; IDO1: cyan; CD11b: green). III: Increased levels of Tregs in and around GME6 (DNA: blue; FOXP3: red; CD3E: green). IV: ICOS positivity primarily were on Tregs (DNA: blue; FOXP3: red; CD3E: green; ICOS: white). V: Lower ICOS positivity in CD8+ T cells (DNA: blue; FOXP3: red; CD8a: green; ICOS: white). VI: Direct interaction between IDO1+ CD11b+ myeloid cells and ICOS+ Tregs denoted by orange arrows (DNA: blue; FOXP3: red; CD11b: green; ICOS: white; IDO1: cyan**). E)** Models showing cellular interaction and relationships in GME6 and GME7.

Comparatively, IDO1^−^ M1 (the most predominant cell type in GME7) had the most proximity to these T cell subtypes in GME7, and several folds lower in GME6. Incidentally, IDO1^−^ DCs had some variability in the volume of proximity to T cell subsets. The spatial proximity volume of IDO1^−^ DCs to CD4^+^ and CD8^+^ T lymphocytes, regulatory T lymphocytes was highest in GME5 (proximity to GZB^+^ CTLs was highest in GME7). These DCs appear to be mostly non-immunosuppressive dendritic cells interacting with T cells in tertiary lymphoid structures (TLS).^37,38^ Collectively, this suggests that the proximity of suppressive DCs to other immune cells likely influences granuloma trajectory towards resolution versus necrosis.

IDO catabolizes tryptophan (Trp) to the product kynurenine (Kyn), resulting in direct and indirect immunosuppression of immune cell populations. IDO1-expressing cells including tolerogenic DCs and macrophages have been reported to have multiple effects on T cells.^39,40^ Previous studies have shown that IDO1 can induce regulatory T cell (Treg) functional activity while inhibiting the proliferation and activation of other T cells.^39–43^ Using ICOS as a marker of activated T cell subsets, we calculated spatial proximity volume to evaluate the interactions between IDO1^+^ DCs and ICOS^+^ and ICOS^−^ T cell subtypes in GME6 and GME7 (**Fig. 6B**).^44,45^ In GME6, IDO1^+^DCs were more closely associated with activated ICOS^+^ Tregs than non-activated ICOS^−^ Tregs. This was the opposite for other T cell subsets, with these suppressive DCs having less interactions with activated T cells (CD4^+^ T cells, GZB^+^ CTLs, other CD8^+^ T cells) than their ICOS^+^ counterparts, while ICOS^+^ CD4^+^ and CD8^+^ T lymphocytes had closer association with IDO1^+^ DCs. Additionally, in GME7, while the trend was the same for CD4^+^ T cells, GZB^+^ CTLs, other CD8^+^ T cells, as in GME6, the spatial volume was near equivalent between ICOS^+^ and ICOS^−^ Tregs. In addition to scoring interaction, it was also important to assess the spatial distance between T cell populations and tolerogenic macrophages and DCs (**Fig. 6C)**. On average, for all T cell subtypes (except Tregs), the ICOS^−^ T cells are closer to IDO1^+^ DCs than the ICOS^+^ populations. However, activated Tregs were more proximal to the IDO1^+^ DCs and macrophages than the non-activated Tregs (**Figs. 6C, D**). Taken together, these data suggest an immunosuppressive program that is related to the proximity and direct interaction between suppressive myeloid cells and activated Tregs, where spatial volume and spatial distance of suppressor cells is associated with diminished activation of nearby T cell subtypes consistent with prior data. This immunoregulatory program is driven by IDO1 positivity in myeloid populations (mainly DCs) (**Fig. 6E**).^12,41,46^

### IDO high ROI are characterized by signaling pathways associated with pathogen control and T cell regulation

Our results show that IDO1^+^ suppressive myeloid cells are differentially present within unique GME. Smaller and more cellular granulomas that appear histologically to be undergoing resolution showed increased numbers of IDO1^+^ suppressor cells subsets. Based on cancer literature, we originally hypothesized that active and necrotizing granulomas would have higher numbers of suppressor cells, yet our data pointed to a protective role of IDO1 in granuloma resolution. Therefore, we wanted to better understand the role of IDO1 in TB infected lung to determine if IDO1-induced transcriptional programs differ in tuberculosis as compared to what has been described in solid tumors. We revisited our spatial transcriptomics dataset, classifying each ROI as either IDO1-high or IDO1-low, irrespective of annotation. We evaluated the genes upregulated in IDO1-high ROI as compared to IDO1-low ROI (**Fig. 7A, Suppl. Table 4**). Genes involved in both suppression (*il1b*, *il17ra*, *s100a8/a9*) and interferon responsiveness (*irf1, stat1, spp1, ifngr2*) were both upregulated in IDO1-high ROI (**Fig. 7B, Suppl. Table 4, Suppl. Fig. 9**). Gene set enrichment analysis showed upregulation of transcriptional programs associated with suppressive and secretory signals including TGF-beta, IL4, and, and IL-10 associated pathways consistent with what has been reported in other disease states.^47–54^ In addition, upregulation of NF-KB and interferon signaling responses and upregulation of antigen presentation was enriched highlighting the complex role of IDO1 in both activating and suppressing immune cell function (**Fig. 7C**). Collectively, transcriptional programs associated with IDO induction mirrors what has been reported in other infectious diseases and cancer.^55–59^

**Figure 7.**
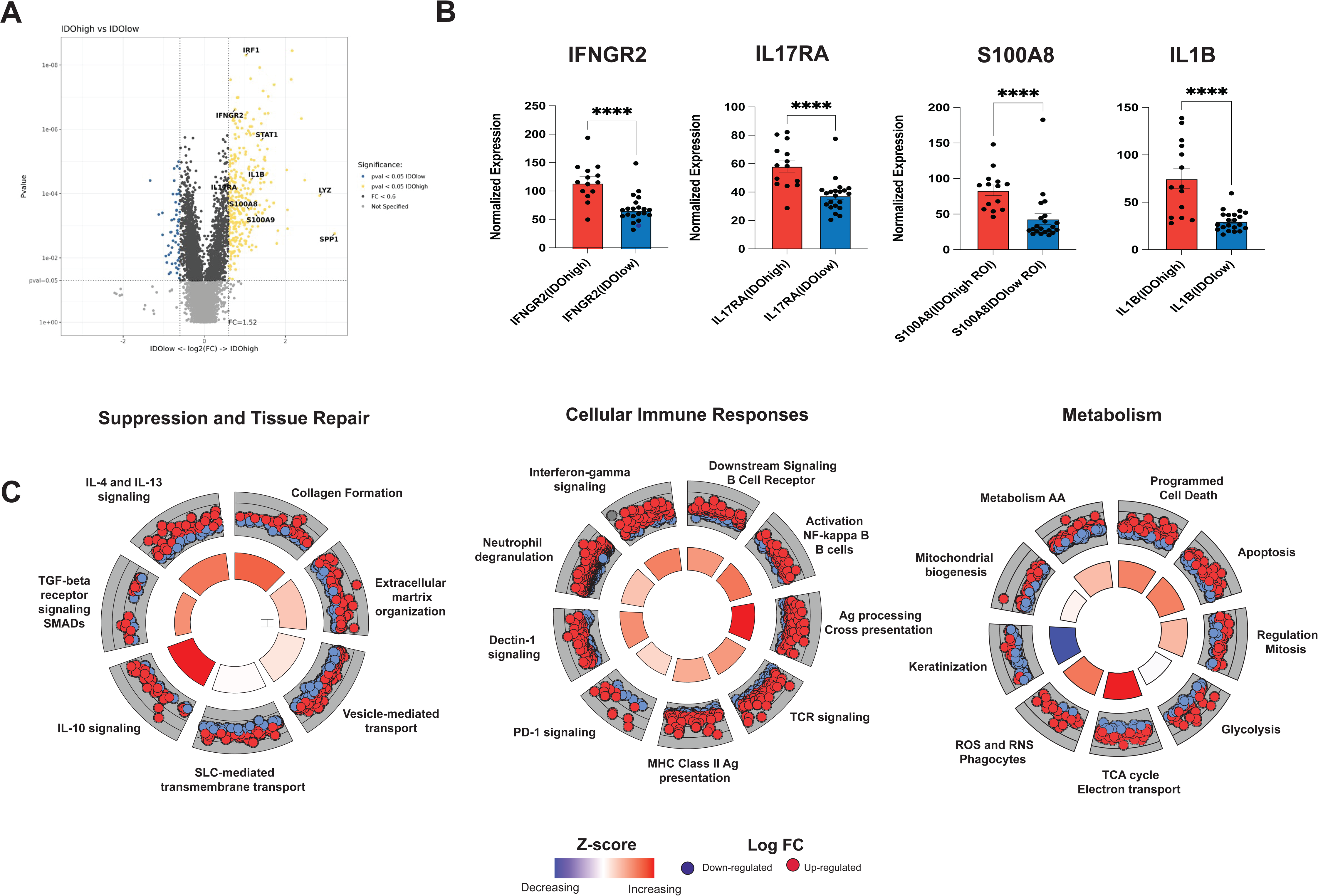
IDO high ROI are characterized by signaling pathways associated with concomitant pathogen control and T cell regulation. **A)** Volcano plot displaying genes significantly upregulated (yellow dots) or downregulated (Blue dots) with greater than 1.5 log fold change on comparing IDO high ROI with IDO low ROI. Some of the important and interesting genes have been marked. **B)** Bar plot of normalized count of 4 genes involved in immune suppression, across the broad classification of IDO high ROI vs IDO low ROI. **C)** GO plot of the broad categories: Secretory and suppression signals, Cellular Immune responses and cell cycle metabolism are highlighted. Each GO plot is comprised of multiple signaling pathways, represented by a pie. Each pathway/pie is represented by genes set [red(upregulated) and blue(downregulated) dots], degree of Normalized enrichment score (NES) and adjusted p value corresponds to color and size of inner circle respectively.

## Discussion

The last decade saw explosion of single cell exploration, spatial technologies and advancement of deeper multiomic understanding of cancer and autoimmunity. But the spatial biology of infectious diseases has been relatively unexplored largely due to limitation in biopsy samples from human patients and ineffective granuloma models in animals that are replicative of human TB granuloma diversity. In addition, TB has a complex pathophysiology, and a limited number of studies have explored cell-cell interactions in TB diseased human lung tissue to interrogate the local immune signatures associated with disease outcome. McCaffery *et al.* showed at single cell resolution, that the spatial organization and co-expression of PDL1, IDO1, and Tregs in the myeloid core are unique features of mycobacterial granulomas in lungs of patients with active disease compared to latent or treated patients. Here we extend those findings by layering transcriptional profile across granulomas with deep immune cell phenotyping in a subset of patients representing reactivation of TB disease to investigate the role of specific suppressive myeloid cell populations in granuloma microenvironments. We profiled a total of 84 granulomas across three TB-infected individuals allowing a detailed investigation of the role of spatial immune interactions on local disease progression. We were able to perform whole slide imaging on these samples and assign phenotypes over 1.9 million cells. Our study shows the remarkable heterogeneity in the cellular composition of pathologically similar granulomas with percentage of macrophages and DCs being notably variable. We identify IDO1^+^ dendritic cells as a main suppressive cell type in non-necrotizing granulomas and investigate how IDO positivity influences local gene expression in TB infected lung reflecting different granuloma maturation states.

MDSCs are considered important cellular immunotherapy targets in solid cancers and autoimmunity. We set out to determine if MDSCs as defined in cancer were present in TB granulomas and associated with the severity of TB active granulomas. MDSCs inhibit function of T lymphocytes — *a priori* we anticipated identifying MDSCs in regions of necrosis due to suboptimal T cell function and decreased macrophage killing of *Mtb*. However, MDSCs are increasingly recognized as a cell state rather than a cell type necessitating the inclusion of both surface and functional markers to define this population in humans/NHPs and mice.

Numerous studies of diseased lung single cell mapping at high resolution have failed to classify MDSCs as a separate entity. This problem is compounded with the usage of limited markers i.e., CD11b^+^Ly6G^+^ cells as PMN-MDSCs or CD11b^+^CD11c^+^ as M-MDSCs resulting in confusion between segregating neutrophils/macrophages from MDSCs. Due to absence of MDSCs in reference data for RNAseq, we could not comment on the percentage of MDSCs across the granulomas or other lung features in our transcriptomics analysis. Using a multiomics approach, we were able to interrogate the presence of suppressive myeloid cell populations, including MDSCs, in TB granulomas. In this study, we decided to quantify MDSCs in human tissue using a combination of surface markers and functional markers reported in the literature. M-MDSCs were identified in both necrotic and cellular granulomas. However, their numbers as compared to other suppressive myeloid populations were extremely low. In addition, we could only detect M-MDSCs and not PMN-MDSCs in TB granulomas. Previous studies studying the role of MDSCs in TB show a high frequency of PMN-MDSCs in peripheral blood associated with active disease and their potential use as a biomarker.^9,20,60^ The presence of PMN-MDSCs in bronchoalveolar lavage (BAL) of pulmonary TB patients has also been reported by multiple studies. Our inability to detect “classically” defined PMN-MDSCs in human lung infected with *Mtb* could be due to these cells only being present in the periphery or that the cells shift their phenotype once migrating into the tissues (*preferred hypothesis).* Alternatively, our inability to detect PMN-MDSCs could potentially be due to the fragile nature of these cells resulting in their absence from FFPE tissues.

Other suppressive myeloid cell populations including IDO^+^ and IDO^−^ dendritic cells and IDO^+^ M1 and M2 macrophages represented the major suppressive myeloid cell populations within TB granulomas in this study. IDO has known roles in immune suppression in cancer and in TB. McCaffrey *et. al.* showed that IDO1^+^ suppressive cells in *Mtb* were seen in active disease TB patients.^12^ D-1MT has been used to neutralize IDO1 in NHP TB. These studies showed that blocking IDO enables granuloma disruption and enhanced TB control in rhesus macaques.^19^ PMN-MDSC were identified in blood and bronchoalveolar lavage (BAL) fluid from macaques infected *M. tuberculosis* CDC1551 and represented major sources of immunosuppressive effectors and mediators (IDO1, IL-10, MMP-9, iNOS, and PD-L1). However, NHP studies represent primary infection, and our analysis is reflective of post-primary or reactivation disease in people. Therefore, traditional MDSCs (M-MDSCs and PMN-MDSCs) could potentially play an important and distinct role during primary infection that could explain dysfunctional anti-TB responses in active TB granulomas.

In coherence with other studies, IDO1 expression in this study was mostly associated with granulomatous regions, but systematic evaluation of 84 granulomas led to differential and significantly upregulated IDO1 protein expression and *ido1* related metabolic pathways within cellular/non-active granulomas in contrast to necrotic/active granulomas. Necrotic granulomas were enriched for neutrophils and IDO1^−^ M1 and M2 macrophages (GME 3 and GME7). Spatial proximity scores and visual inspection of the organization of the GME in necrotic granulomas indicate a more diffuse arrangement of cells across GME containing resting IDO1^−^ myeloid cells as compared to cellular granulomas. Cellular granulomas have significantly “tighter” association with suppressive myeloid cells and both regulatory and traditional T cells indicative of contact-dependent granuloma remodeling associated with high IDO expression in myeloid cells. The most abundant suppressive myeloid cells identified in TB granulomas were suppressive dendritic cells defined as IDO1^+^ CD11b^+^ CD11c^+^. IDO1^+^ DCs were enriched in a microenvironment that included many other immunosuppressive myeloid cells including IDO1^+^ M1, IDO1^+^ M2 macrophages, and in addition to a number of M-MDSCs which were almost all IDO^+^ and PD-L1^+^. Our finding that cellular granulomas have more IDO1^+^ DCs that closely associate with other IDO1^+^ suppressive myeloid cells and T-cells in the context of diminished tissue pathology (lack of necrosis) suggests that suppressive signaling is essential to tissue repair, remodeling, and granuloma resolution. Therefore, suppressive myeloid cells may play a critical role in reducing lung pathology, and our data suggests that further investigation into a potential “protective” role of suppressive myeloid cells in TB is necessary to understand whether HDT targeting suppressive cells is warranted in TB. We propose a model by which granuloma formation is driven by the induction of suppressor signals in the lymphocytic cuff, yet the fate of the granuloma towards resolution versus progression (necrosis) is dictated by the effective transition of myeloid cells to an increasingly suppressive state characterized by closer and tighter cell-cell association of suppressive or tolerogenic DCs with T lymphocyte populations. In this model, suppressor signaling is protective and essential for granuloma resolution. The metabolic transition of myeloid cells to a tolerogenic state may be necessary to induce a concomitant metabolic reprogramming of intracellular *Mtb* to enter a dormant state furthering granuloma resolution.

Limitations of this study include a small sample size of TB patients that underwent pulmonary resection for active disease or biopsy to rule out neoplastic pulmonary disease which means this patient subset represents patients with post-primary rather than primary TB disease. Yet across the three patients, we were able to profile a total of 84 granulomas of varying histological classification allowing a detailed investigation of the role of spatial immune interactions on local disease progression. Importantly, we demonstrate the benefit and necessity of multimodal tissue profiling utilizing both spatial transcriptomics and single-cell immunophenotyping with CyCIF to adequately define and phenotype suppressor cell populations in TB infected lung. While single-cell segmentation in tissue specimens can sometimes incompletely resolve phenotypes of cells in direct contact due to lateral spillover of cell surface markers, this limitation is generally overcome by the ability to profile large areas of tissue, appropriate image quality control, and utilizing a very systematic phenotyping workflow (**Suppl. Fig. 5**). Further studies profiling increased numbers of human granulomas will help determine if the role of suppressor cells in orchestrating individual granuloma trajectories is consistent in primary and post-primary TB disease.^12,61–63^

It has long been appreciated that *Mtb* has evolutionarily conserved T cell epitopes in its genome. This has suggested that T cell recognition of *Mtb* infected macrophages is important for *Mtb* pathogenesis but exactly how or why this may be occurring has been understudied.^64^ Our data supports a model by which T cell recognition of infected macrophage promotes macrophage killing of *Mtb* but also necrosis which leads to lesion expansion, spread, and potential rupture into airways that promotes *Mtb* transmission. It also points to the role of suppressive myeloid cells in tamping down inflammation associated with *Mtb* infection, promoting *Mtb* persistence in the lung without lung damage due to decreased T cell activation even in the presence of antigen presentation and T cell recognition.

Our data highlights the critical role of IDO in TB pathology and granuloma maturation. Intracellular pathogens such as *Chlamydia, Toxoplasma gondii* amongst others require host tryptophan to survive, making IDO1-mediated depletion of intracellular tryptophan levels, in concert with interferon-gamma secretion, a key host defense, starving the pathogens of essential tryptophan.^65–68^ It has been previously shown that tryptophan synthetic machinery of tuberculosis is required to counter CD4^+^ T cell pressure in vivo. Zhang *et.al.* showed that *Mtb* tryptophan auxotrophs are preferentially cleared from TB infected mice.^66^ IDO neutralization has been proposed as a potential adjunct host-directed therapy for *Mtb* with the intent to decrease MDSCs and provide “immune resuscitation” for T cells within granulomas to promote killing of *Mtb* infected macrophages.^11^ However, our data suggests that therapeutics aimed at decreasing suppressive myeloid cells may actually induce *Mtb* disease and pathogenesis depending on the timing of administration. In our model disrupting organized granuloma structure does not enhance T cell mediated killing of *Mtb* but rather the opposite with over-exuberant inflammation leading to ineffective T cell function and lesion progression. It is possible that host initiated suppressive signaling influences mycobacterial metabolic state, transition of *Mtb* to dormancy, and more effective clearance of *Mtb* within granulomas. Future studies evaluating the effect of suppressor cell targeted therapeutics are needed to understand the temporal role of suppressive myeloid cells on lesion development during primary versus post-primary disease to better understand therapeutic windows for drugs targeting these cells. It is also likely that the role of suppressor cells shifts throughout the course of infection.

Future studies using animal models including non-human primate models provide an opportunity to study the role of suppressor cells on early granuloma formation, granuloma maturation, and granuloma resolution in the context of drug treatment.

## Author Contributions

NJ, EO, and AJM wrote the manuscript with all the authors. LZ, ZM, EO, CJ, NJ, and HK performed the experiments. AS, AG, JMR, and IS performed pathological and clinical assessments. SS, PKS, BA, and AJM designed and supervised the experiments.

## Acknowledgments

We would like to acknowledge funding from Gates Foundation grant INV-027106 (PKS), Harvard University Center for AIDS Research P30 AI060354 (JMR), Harvard Clinical and Translational Science Center K12 TR002542 (JMR), and NIAID R21 AI155003-01 (PKS, BA, AJM).

## Declaration of interests

JMR reports compensation for consulting from Third Rock Ventures, and he began employment with Merck after his contributions to this manuscript.

**Suppl. Table 1.** Clinical metadata TB cases.

**Suppl. Table 2.** Complete list of DEG across annotations and list of unique DEG in each annotation.

**Suppl. Table 3.** t-CyCIF TB antibody panel.

**Suppl. Table 4.** Complete list of DEGs for IDO1 high ROI versus IDO1 low ROI.

**Suppl. Fig. 1. Fluorescent protein expression across ROI selected for GeoMx profiling**; CD3, CD68/CD163, and vimentin.

**Suppl. Fig. 2. GeoMx QC and normalization.** Q3 normalization, number of cell nuclei per ROI, and normalized gene counts for CD3 and CD68 across selected ROI highlighting ROI that are T-cell rich versus macrophage rich.

**Suppl. Fig.3. Proportion of cell-types by cell deconvolution (see also Fig.2).**

**Suppl. Fig. 4. Heatmap of DEG across annotations.** Unsupervised clustering of DEGs across annotations (CG, NC, LN and TLS) in comparison to uninvolved lung (UL)s.

**Suppl. Fig. 5. Main phenotyping workflow for assigning cell types**. Unknown or unassigned cells are excluded from subsequent spatial analysis.

**Suppl. Fig. 6. A)** Exemplary CyCIF images showing M-MDSCs in TB granulomas: I: M-MDSCs phenotyped as CD11b+CD33+HLA-DRB1-CD14+ cells. II: IDO1+ M-MDSCs. III: PD-L1+ M-MDSCs. M-MDSCs are denoted by orange arrows. **B)** Proportion of cell phenotypes in necrotic and cellular (non-necrotic) granulomas.

**Suppl. Fig. 7. A**) Spatial scatter plot of whole-slide tissue samples, color-coded by spatial microenvironment (left) and phenotype (right). **B, C).** Heatmap and box plots showing the shortest distance between different GMEs especially GME6 and GME7.

**Suppl Figure 8**. **A. Spatial analysis revealing interaction between cell types quantified across all the images.** This assesses significant co-localization not due to random chance. Here, a k-nearest neighbor method was also used with k=10 to define neighbors immediately adjacent to each cell. A large component of the immunosuppressive GME6 comprised of IDO1+ DCs. IDO1+ DCs had significant interactions with Tregs, other CD4 T cells, GZB+ CTLs, other CD8 T cells, IDO1+ M1 and M2 macrophages.

**Suppl. Fig. 9. Additional significant and non-significant genes in IDO1 hi versus IDO1 lo.**

## Methods

### Patients

3 Archived human FFPE tissue were selected from the tissue repository of MGH and BWH based on positive acid fast *Mtb* staining, culture, or PCR confirming MTB complex infections, and variations in granulomatous inflammation and T lymphocyte infiltration as scored by a pathologist. Specimens were collected under MGH IRB 2019P001114 and BWH IRB 2018P001627.

### Sample Acquisition (GeoMx)

5uM serial sections were freshly cut for GeoMx ^13,69^. Consecutive serial section was used for H&E staining, CD4 IHC, CD8 IHC, Acid-fast staining and GeoMx. GeoMx slides were processed for deparaffinization and antigen-retrieval based on GeoMx guidelines. Briefly, we used the whole transcriptome atlas (WTA) RNA panel to understand the TB spatial biology. We used in-house morphology/visualization antibody markers (SYTO13(Blue):DNA, AF594:CD68(clone KP1, Santa Cruz, sc-20060AF594)/CD163(clone EPR14643-36, Abcam ab272103), AF647(red):CD3E(Clone UMAB54, Origene, UM500048), AF488(green):Vimentin(Clone E5, Santa Cruz, sc-373717), in combination with H&E staining and pathologist opinion to select the Region of interest (ROI) to include range of pathology features in diseased lung. In total, 36 ROIs were marked form all 3 cases, 6 uninvolved regions, 5 necrotic granulomas, 15 cellular/non-necrotic granulomas, 4 Tertiary lymphoid structure and 5 pulmonary associated Lymph nodes.

After ROIs selection, we collected the spatial RNA pattern information as Indexed oligos from each ROI to specific well of an Illumina microplate. Subsequently, Indexed oligos are hybridized to GeoMx hyb codes and analyzed using Illumina NovaSeq6000 to generate digital counts of the RNA.

The raw counts of RNA seq data was obtained from Illumina NovaSeq6000. Quality control was performed in GeoMx DSP software version 3.1. Parallelly we also analyzed data using DEseq2 (NGS pipeline) which has an inbuilt QC analysis module. All QC data was normalized using Q3 normalization method. The normalized data matrix was then used for all downstream analysis. Different dimensionality reduction algorithms were performed: Principal component analysis (PCA), UMAP and tSNE using DSP software R module. Cluster analysis (Unsupervised and Supervised) were performed using DSP software R module. Unsupervised clustering of all ROI led to selection of 3 uninvolved lung which clustered together as a reference for comparative analysis.

Image of serial sections and region of interest were selected by GeoMx DSP software version 3.1. Photocleavable oligonucleotide tags were hybridized to targets on slide mounted sections. Oligonucleotide tags were released via UV exposure and the sequencing library was prepared using an Illumina NextSeq 2000 P3 kit. Pooled library was sequenced on Illumina NovaSeq 6000 instrument. FASTQ files were processed by GeoMx NGS Pipeline v3.1 to remove low quality reads and adapter sequences. Read count matrix was obtained by aligning FASTQ reads to oligonucleotide tags specific for each gene. Differential gene expression analysis was executed by the DESeq2 pipeline. Briefly, raw count was normalized for differences in library depth and gene-wise dispersions was estimated. A negative binomial model was fit and hypothesis testing was performed using likelihood ratio test to estimate differential expressed genes between conditions. For gene-set enrichment analysis fgsea R package (fast pre-ranked gene set enrichment analysis) was used with minSize = 10, maxSize = 500 and nperm = 10000 parameters. Gene set “c5.go.bp.v7.4.symbols.gmt” was downloaded from MSigDB database and used in analysis.

### t-CyCIF Protocol and Image Acquisition

Tissue-based cyclic immunofluorescence (t-CyCIF) was performed on formalin-fixed, paraffin-embedded (FFPE) tissue sections, adapting established protocols (https://dx.doi.org/10.17504/protocols.io.5qpvorbndv4o/v2)^14,70,71^ The BOND RX Automated IHC/ISH Stainer was used to pre-process the slides, which were first baked at 60°C for 30 minutes, followed by dewaxing with Bond Dewax solution at 72°C. Antigen retrieval was carried out using Epitope Retrieval 1 solution (Leica™) at 100°C for 20 minutes. The samples then underwent several rounds of antibody incubation, imaging, and fluorophore inactivation. All primary and secondary antibodies used during CyCIF study are listed in **Supplemental Table 2**. Antibodies were incubated in the dark overnight at 4°C, with Hoechst 33342 used for nuclear staining during each cycle. After wash steps, coverslips were wet-mounted using 70% glycerol in Phosphate-buffered saline (PBS). Images were acquired on a CyteFinder slide scanning fluorescence microscope (RareCyte Inc.), using a 20x/0.75 NA objective – with a 2×2 binning. Fluorophore inactivation was achieved using a 4.5% hydrogen peroxide (H₂O₂) and 24 mM sodium hydroxide (NaOH) solution under LED illumination for an hour.

### Image Processing and Quality Control

Images were processed using the Docker-based NextFlow pipeline MCMICRO (https://github.com/labsyspharm/mcmicro).^72^ First, illumination correction was performed with flat fielding using the BaSiC software.^73^ For each sample, all the raw images from individual cycles were stitched and registered into an aligned mosaic multiplexed image using the ASHLAR module.^74^ The assembled images were next prepared for single-cell quantification by the generation of probability maps using the UnMicst v1 model ^75^ and single-cell segmentation by the marker-controlled watershed S3segmenter (https://github.com/HMS-IDAC/S3segmenter). Background subtraction was applied using an ImageJ/Fiji rolling ball algorithm with 50-pixel radius (https://github.com/Yu-AnChen/imagej-rolling-ball). Following the generation of segmentation masks, MCQuant extracted the fluorescence intensity of each marker for individual cells – producing a single-cell data table for the whole-slide image. To ensure the quality of the single-cell data, the staining of each marker was reviewed and evaluated in all the samples before being used for further analysis. Additionally, the interactive QC software, CyLinter, was used to remove regions containing tissue and staining artifacts and filter out cells with no or dim nuclei, cells lost in the multi-cycle experiment, and cells with outlier marker staining.^76^

### Multiplexed Image Analysis

All the markers used for downstream analysis were individually thresholded using an open-source visual gating tool (https://github.com/labsyspharm/minerva_analysis). Single-cell image analysis was performed using the Python package SCIMAP.^77^ The obtained gates for each marker were used to rescale the single-cell image data, scaling it between 0 and 1 (with values above 0.5 indicating marker positivity). Cell-type calling was defined based on gating strategies adapted from flow cytometry panels used for identifying multiple immune and non-immune cell types (**Supplemental Figure 3; Supplemental Table 3**). Markers used for phenotyping include aSMA, panCK, CD31, CD45, CD206, CD11b, CD33, CD14, CD66b, HLA-DRB1, CD20, CD3E, CD4, CD8a, FOXP3, CD163, CD11c, MCT, CD16, GZB, CD68, IDO1 (**Supplemental Table 2**).

Spatially recurrent cell neighborhoods were defined by SCIMAP function (*scimap.tl.spatial_lda*). Neighborhood matrices were based on spatial phenotypic patterns using the Latent Dirichlet Allocation algorithm to identify RCNs. LDA is an unsupervised generative probabilistic learning model that is widely utilized to organize large datasets so as to uncover insights into the underlying thematic data structure ^78^. In this instance, the algorithm modeled the latent space of cellular distributions and identified spatially-organized recurrent cellular neighborhoods (RCNs) – also known as microenvironments. In essence, we trained the models using a k-nearest neighbor (kNN) method for the cellular neighborhood definition, identifying neighbors as within a 10-cell proximity of every cell. The latent weights or variables generated post-training were extracted and then clustered using k-means clustering. The RCNs were clustered into 8 initial environments. Two similar epithelia-rich MEs were merged to yield ME1, resulting in a total of 7 microenvironments. Other spatial analyses were performed using SCIMAP functions including spatial distance (*scimap*.*tl.spatial_distance*), spatial interaction (*scimap.tl.spatial_interaction*) and proximity volume (*scimap*.*tl.spatial_pscore*, https://scimap.xyz).^77^

