## Supplementary figures and images for "Multiomic analysis identifies suppressive myeloid cell populations in human TB granulomas"

### Supplemental Figure 1

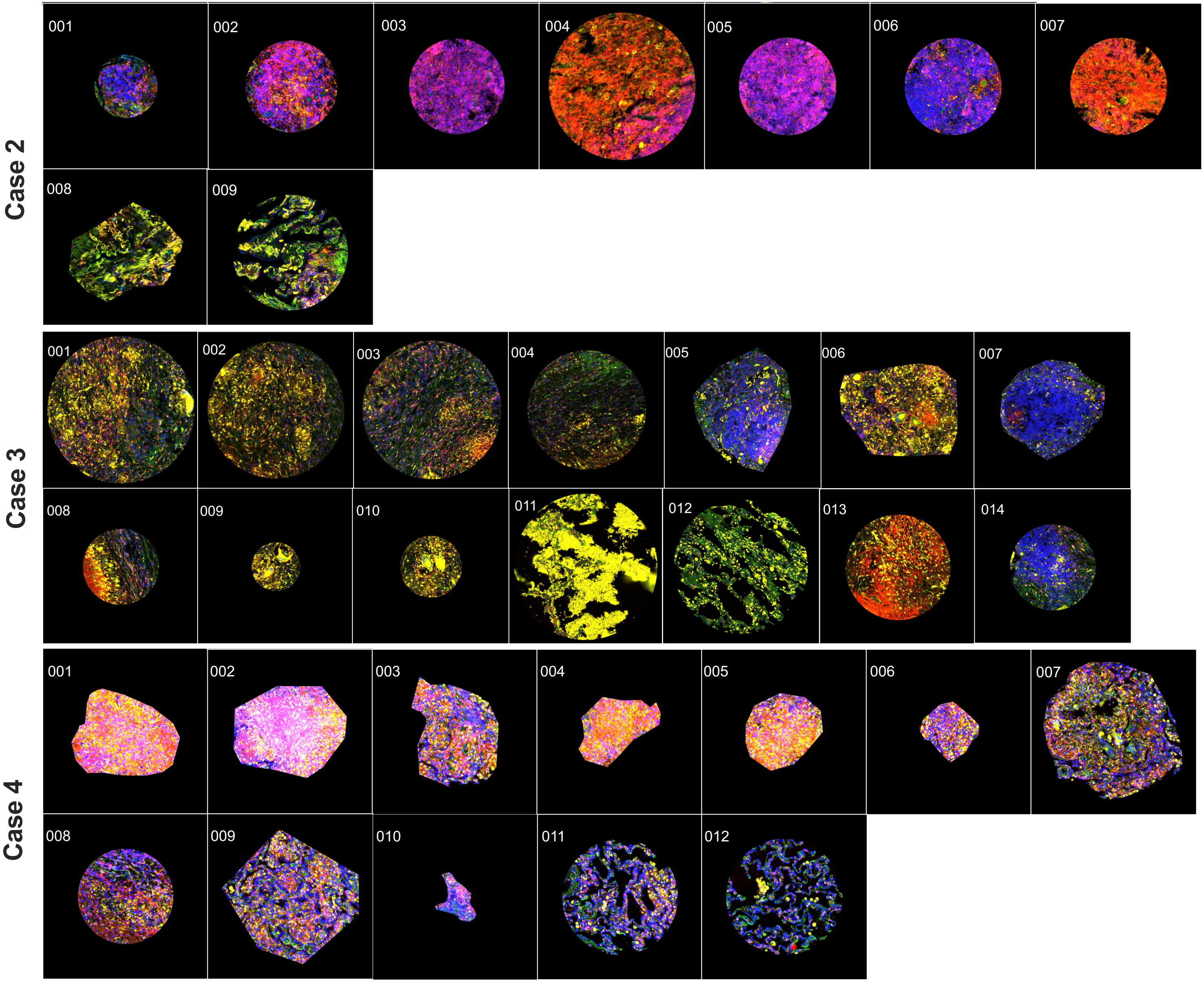

### Supplemental Figure 2

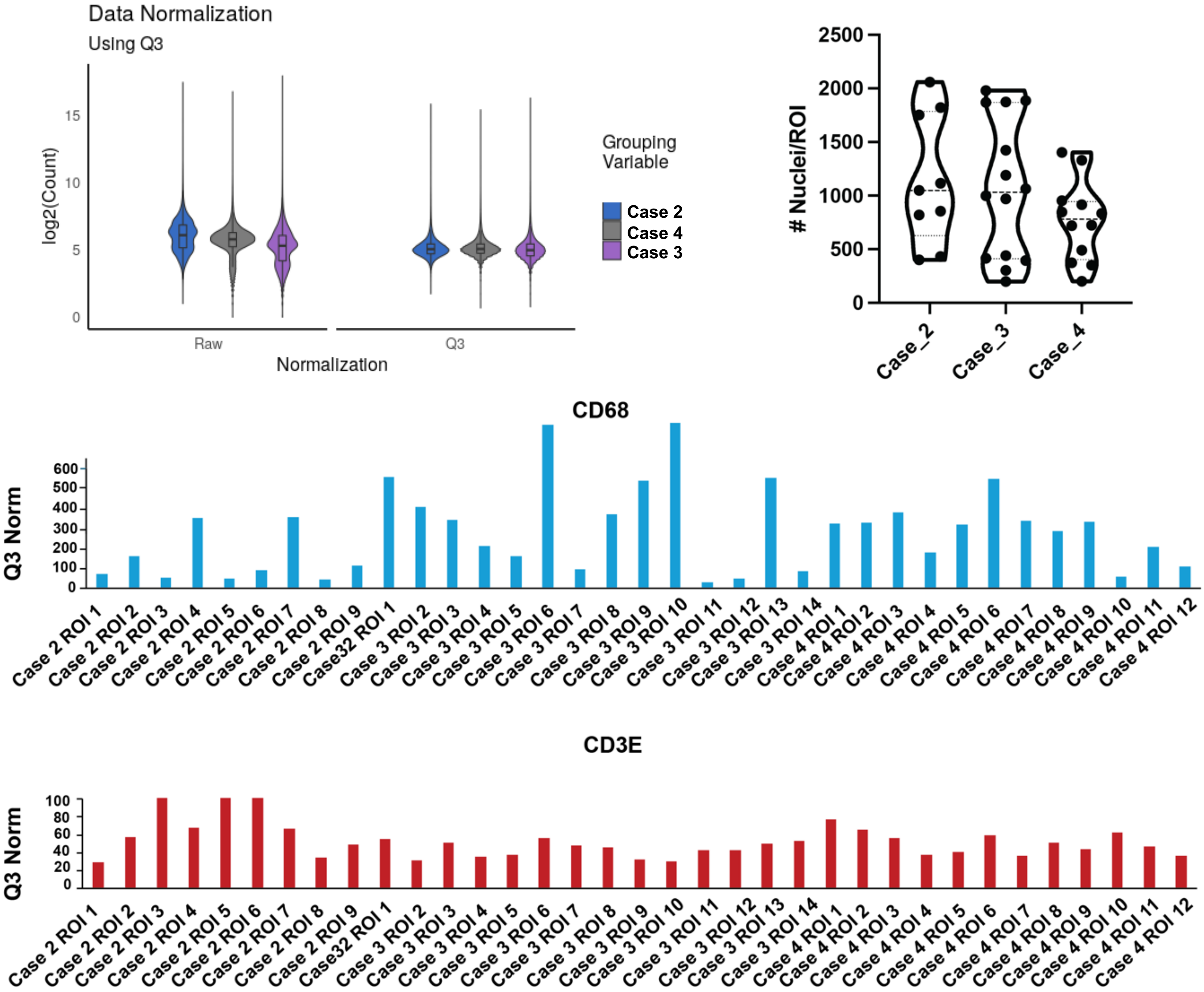

### Supplemental Figure 3

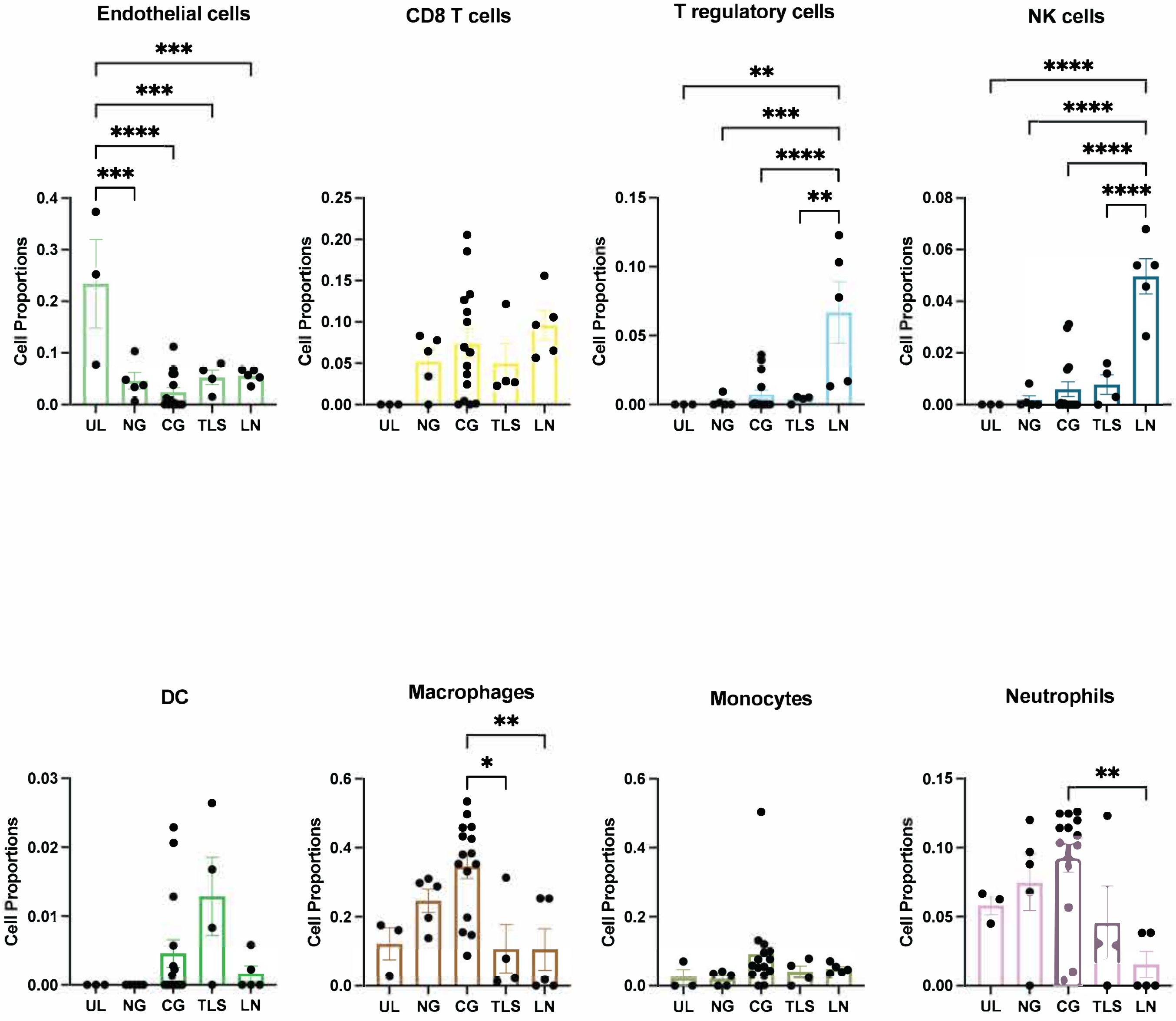

### Supplemental Figure 4

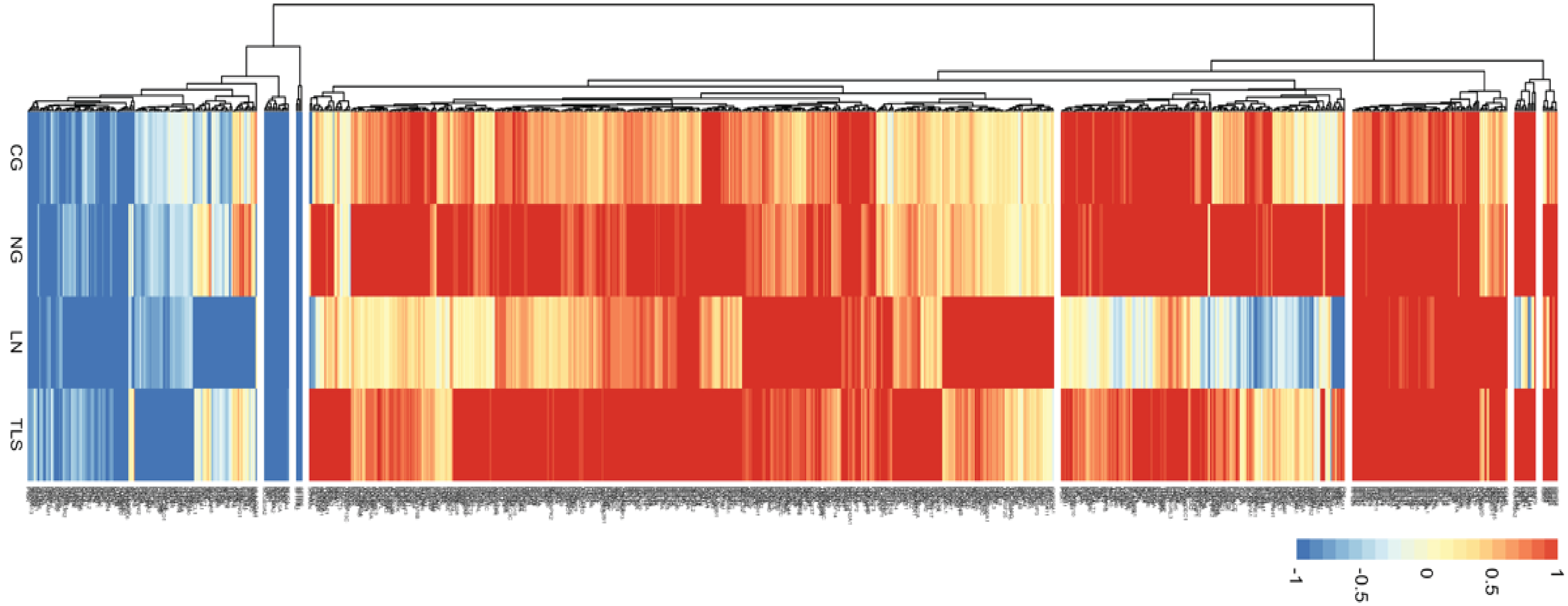


**Suppl. Figure 4**

### Supplemental Figure 5

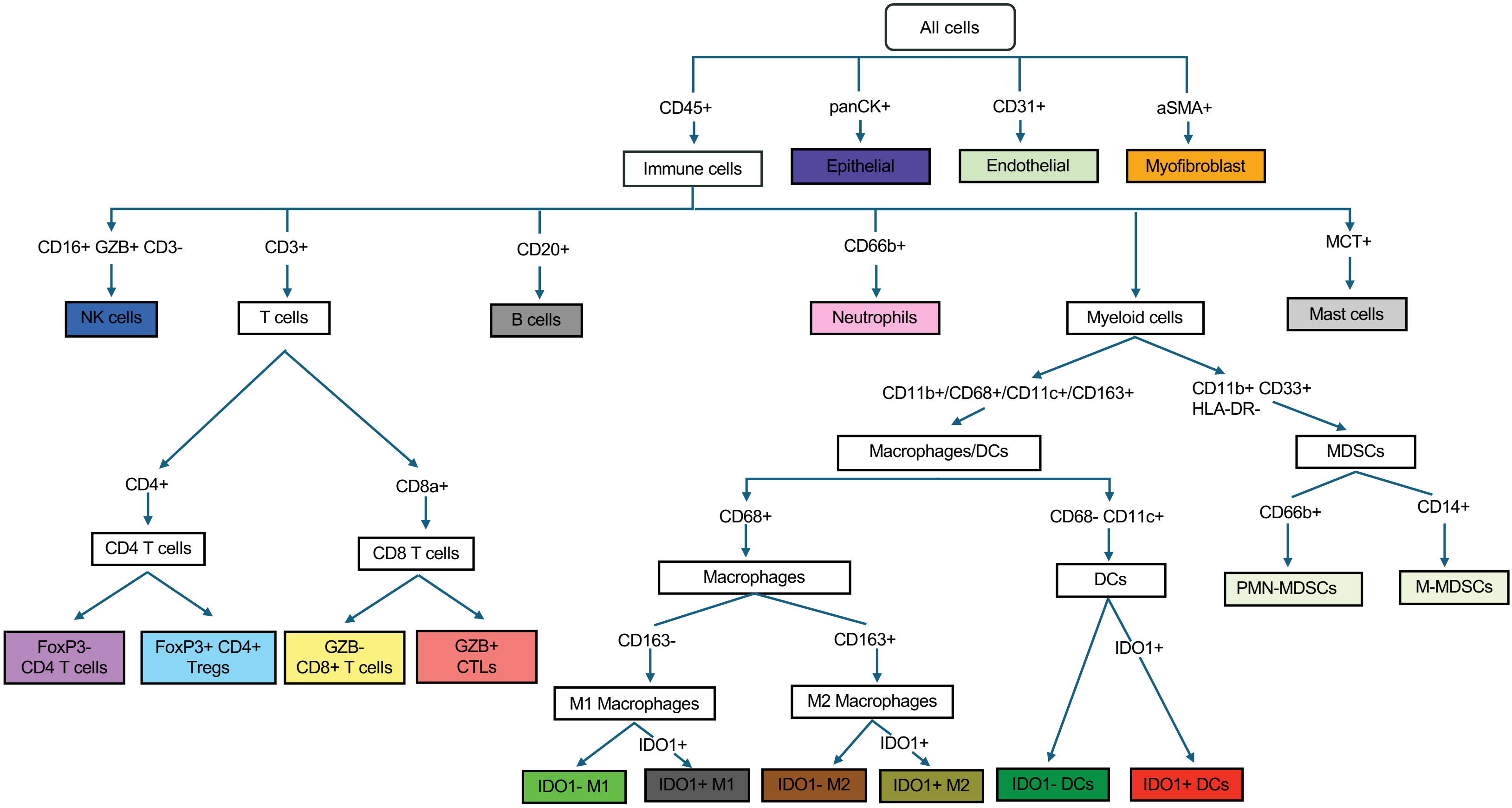

### Supplemental Figure 6

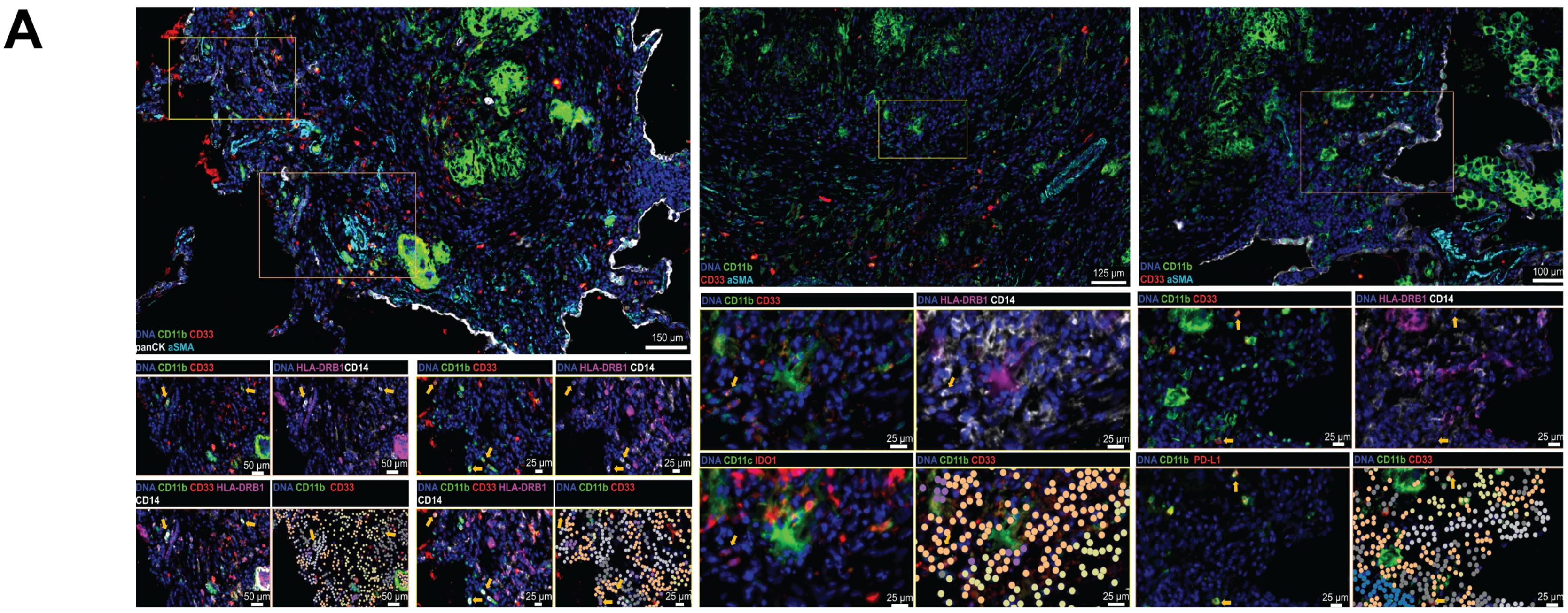

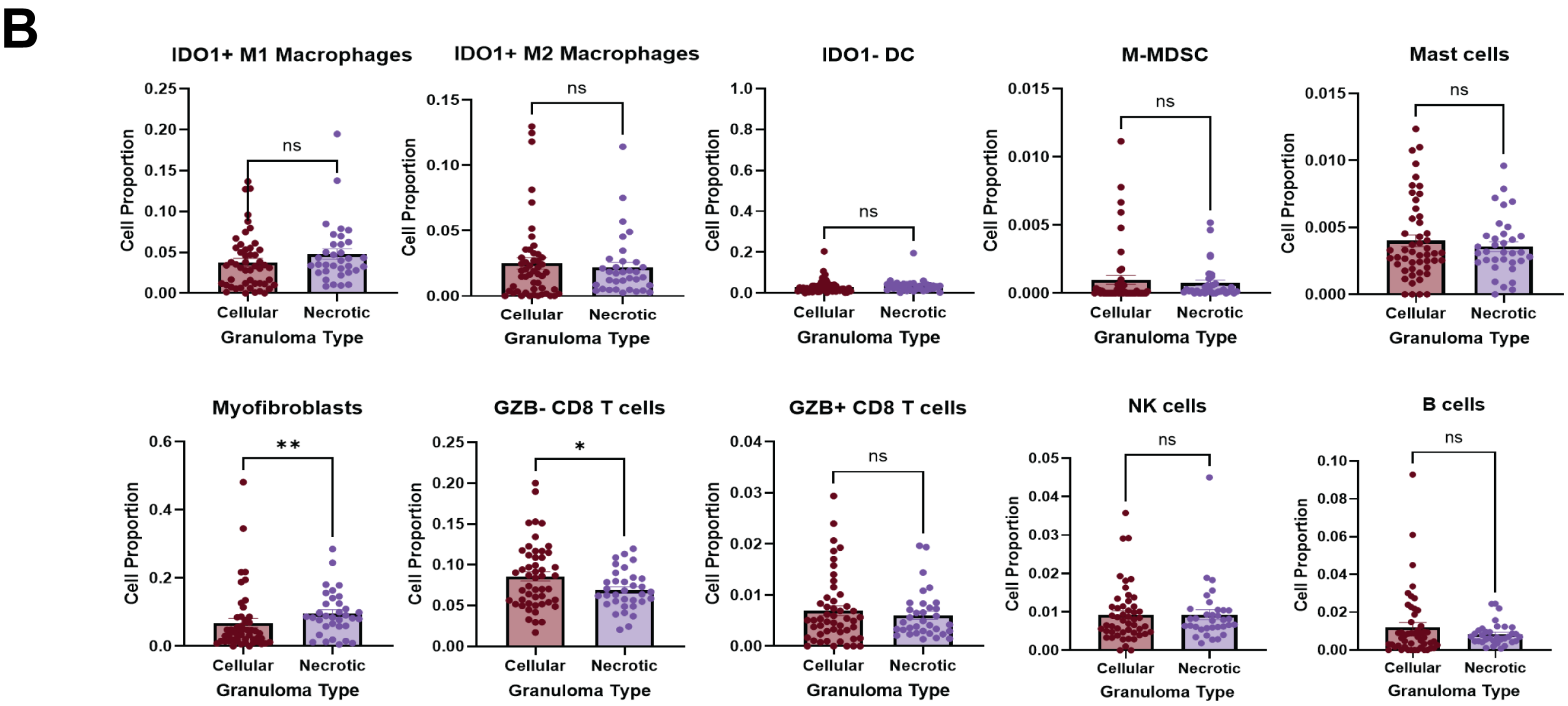

### Supplemental Figure 7

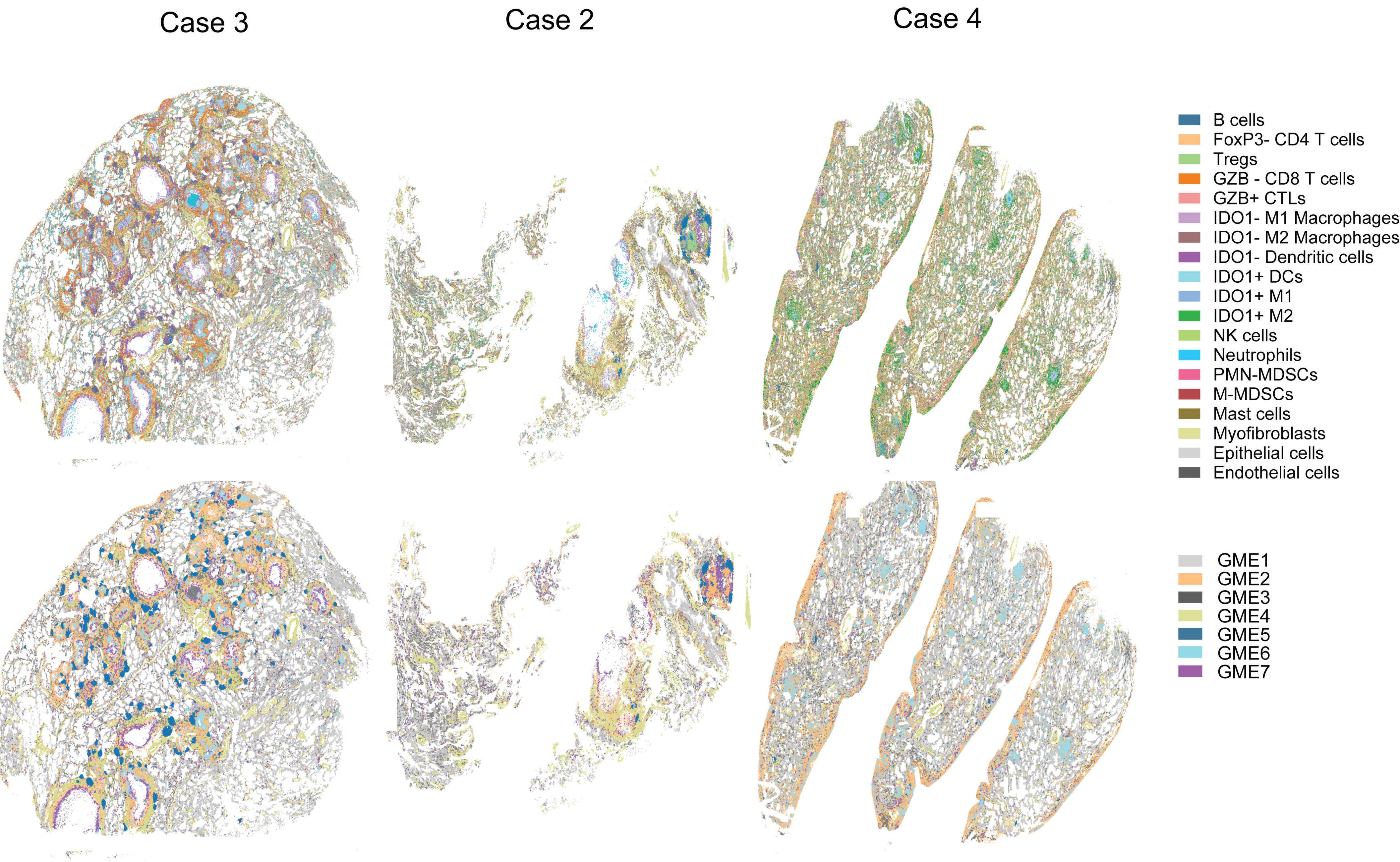

### Supplemental Figure 8

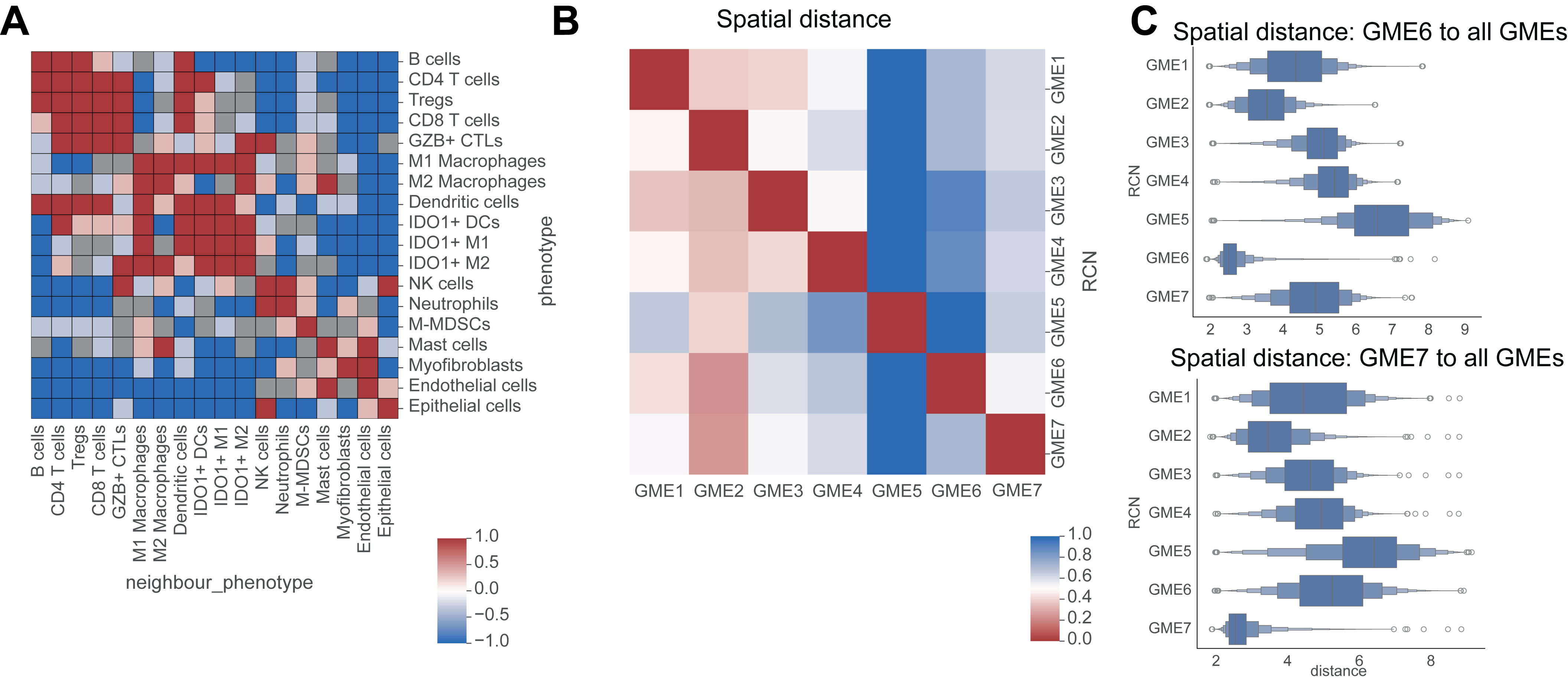

### Supplemental Figure 9

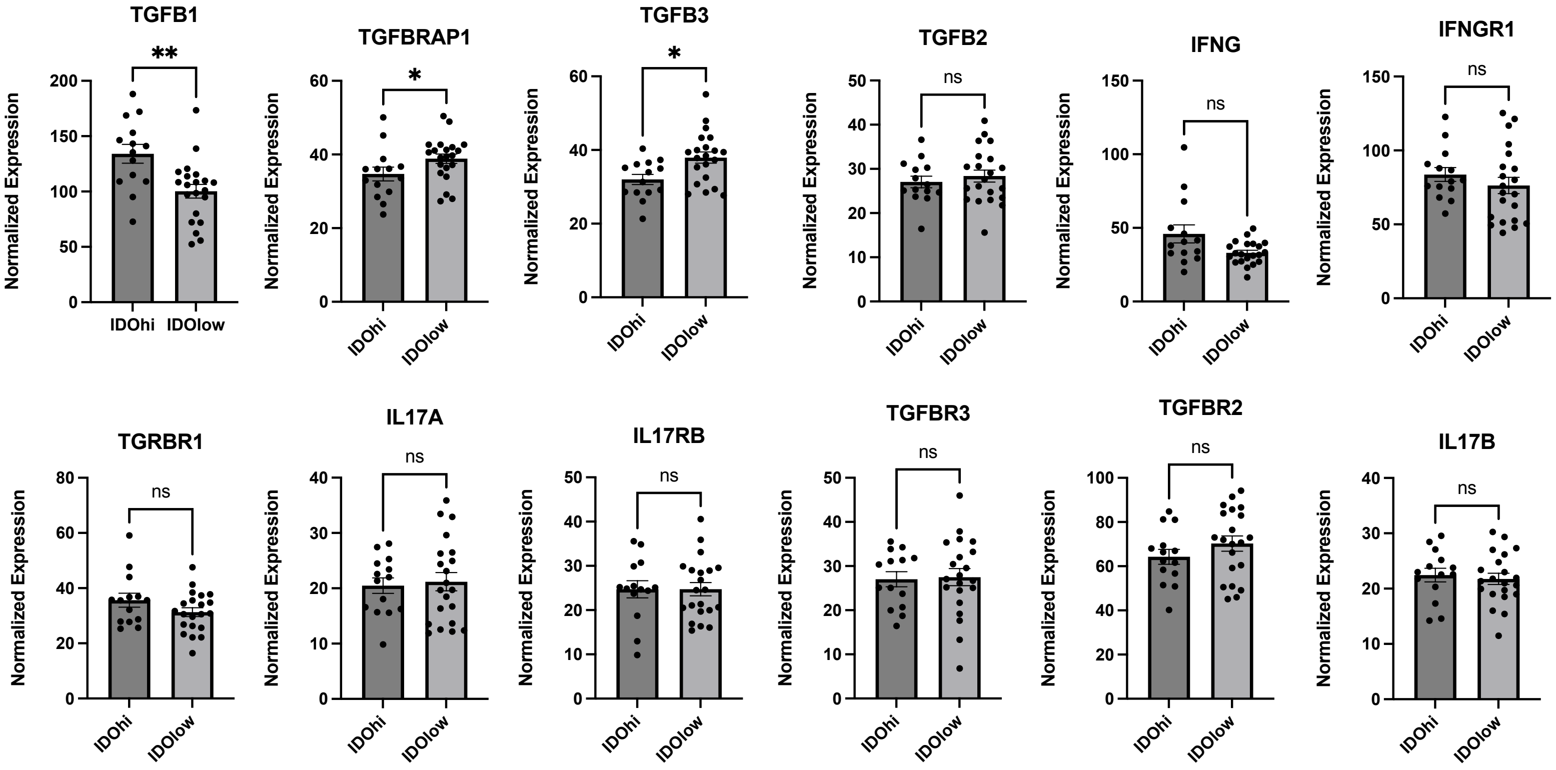
